# Systemic Tissue and Cellular Disruption from SARS-CoV-2 Infection revealed in COVID-19 Autopsies and Spatial Omics Tissue Maps

**DOI:** 10.1101/2021.03.08.434433

**Authors:** Jiwoon Park, Jonathan Foox, Tyler Hether, David Danko, Sarah Warren, Youngmi Kim, Jason Reeves, Daniel J. Butler, Christopher Mozsary, Joel Rosiene, Alon Shaiber, Ebrahim Afshinnekoo, Matthew MacKay, Yaron Bram, Vasuretha Chandar, Heather Geiger, Arryn Craney, Priya Velu, Ari M. Melnick, Iman Hajirasouliha, Afshin Beheshti, Deanne Taylor, Amanda Saravia-Butler, Urminder Singh, Eve Syrkin Wurtele, Jonathan Schisler, Samantha Fennessey, André Corvelo, Michael C. Zody, Soren Germer, Steven Salvatore, Shawn Levy, Shixiu Wu, Nicholas Tatonetti, Sagi Shapira, Mirella Salvatore, Massimo Loda, Lars F. Westblade, Melissa Cushing, Hanna Rennert, Alison J. Kriegel, Olivier Elemento, Marcin Imielinski, Alain C. Borczuk, Cem Meydan, Robert E. Schwartz, Christopher E. Mason

## Abstract

The Severe Acute Respiratory Syndrome Coronavirus 2 (SARS-CoV-2) virus has infected over 115 million people and caused over 2.5 million deaths worldwide. Yet, the molecular mechanisms underlying the clinical manifestations of COVID-19, as well as what distinguishes them from common seasonal influenza virus and other lung injury states such as Acute Respiratory Distress Syndrome (ARDS), remains poorly understood. To address these challenges, we combined transcriptional profiling of 646 clinical nasopharyngeal swabs and 39 patient autopsy tissues, matched with spatial protein and expression profiling (GeoMx) across 357 tissue sections. These results define both body-wide and tissue-specific (heart, liver, lung, kidney, and lymph nodes) damage wrought by the SARS-CoV-2 infection, evident as a function of varying viral load (high vs. low) during the course of infection and specific, transcriptional dysregulation in splicing isoforms, T cell receptor expression, and cellular expression states. In particular, cardiac and lung tissues revealed the largest degree of splicing isoform switching and cell expression state loss. Overall, these findings reveal a systemic disruption of cellular and transcriptional pathways from COVID-19 across all tissues, which can inform subsequent studies to combat the mortality of COVID-19, as well to better understand the molecular dynamics of lethal SARS-CoV-2 infection and other viruses.

## Introduction

In March 2020, the World Health Organization (WHO) declared a novel pandemic of the coronavirus disease 2019 (COVID-19), an infection caused by the betacoronavirus Severe Acute Respiratory Syndrome Coronavirus 2 (SARS-CoV-2)^1^, currently attributed to over 115 million cases and over 2.5 million deaths globally (https://coronavirus.jhu.edu). Since the presenting symptoms of COVID-19 resemble those of common viral respiratory infections, a molecular diagnosis is required to distinguish a SARS-CoV-2 infection from influenza and other respiratory illnesses^2,3^, and ongoing questions remain about the host responses to SARS-CoV-2 relative to other respiratory pathogens. As severe illness and death continue to impact a segment of COVID-19 positive individuals, urgent questions about the molecular drivers of morbidity and mortality associated with SARS-CoV-2 infection persist. This knowledge could lead to improvements in both the acute treatment and long-term management of pathological changes in multiple organs.

Prior work has shown that COVID-19 creates systemic and severe interferon response (both alpha and gamma), and that co-infection with other pathogens is relatively rare (3-15%)^4,5,6^. Yet, there is limited data for discriminating the molecular response between different kinds of respiratory infections or pulmonary conditions (e.g. influenza vs. COVID-19) and almost no data on the variegated impact of different pathogens across different tissues. Delineation of pathogen- and tissue-specific differences is critical for understanding the molecular determinants of mortality associated with COVID-19, as well as guide development of novel diagnostics and therapeutic interventions.

To address these gaps in knowledge, we first used shotgun metatranscriptomics (total RNA-seq) to comprehensively profile human tissues in 39 patients who died from COVID-19 (n=190 total autopsy samples), including heart, liver, lung, kidney, and lymph nodes, and analyzed gene expression, isoform, and T cell receptor expressed motifs (TCEMs) alterations. We also used a spatial protein and transcript mapping platform (GeoMx) to delineate the cartography of the infection in these tissues and to discover disruptions of cell-to-cell interactions. The spatial omics data examined 357 areas of interest (AOIs) from 16 total patients with COVID-19, influenza, ARDS, and normal, revealing the cellular and regulatory signatures that define these distinct pathological states. Finally, to provide context to earlier stages and sites of infection, we compared these in-depth spatial and tissue-specific transcriptome maps with an independent cohort of nasopharyngeal (NP) swabs from 216 COVID-19 positive patients and 430 COVID-19 negative controls, which reveal a significant, distinct disruption of cellular and transcriptional programs induced by SARS-CoV-2 infection in the patients who unfortunately succumbed to the disease. These data were also placed into an online portal for data mining and visualization at https://covidgenes.weill.cornell.edu/.

## Results

### Spatial and expression profiling of high and low SARS-CoV-2 infection

We first used the GeoMx Digital Spatial Profiling (DSP) platform to perform multiplexed high-resolution spatial transcriptomic profiling of 357 lung tissue regions of interest (ROIs) from 16 patients. These were selected from deceased patients with COVID-19 (n=31), non-viral ARDS (2), influenza induced ARDS (3), and healthy tissues from individuals without infections as controls (3) using nCounter Multiplex Analysis which incorporated targets for SARS-CoV-2 (i.e. COVID-19 Spike-In) (See Methods, **Table 1**). Among 31 COVID-19 donors, we identified lung samples that had high overall SARS-CoV-2 expression (COVID-19 high) or had low overall expression of SARS-CoV-2 (COVID-19 low), and four representative samples from each group were selected for downstream analysis (**Extended Data Figure 1**). Serial sections were stained with an RNAscope probe against the viral S gene, Syto13 (nuclear DNA), Macrophages (CD68), immune cells (CD45), and epithelial (Pan-cytokeratin) along with the GeoMx Cancer Transcriptome Atlas Panel (CTA, 1811 targets), supplemented with 23 human genes associated with lung biology and two ORFs from the SARS-CoV-2 genome (See **Methods**, **Extended Data Figure 1**). We chose tissue regions (Regions of Interest, ROIs) that captured three structural components of the lung, including vascular, airway, and alveolar regions (**Figure 1a**).

**Figure 1.**
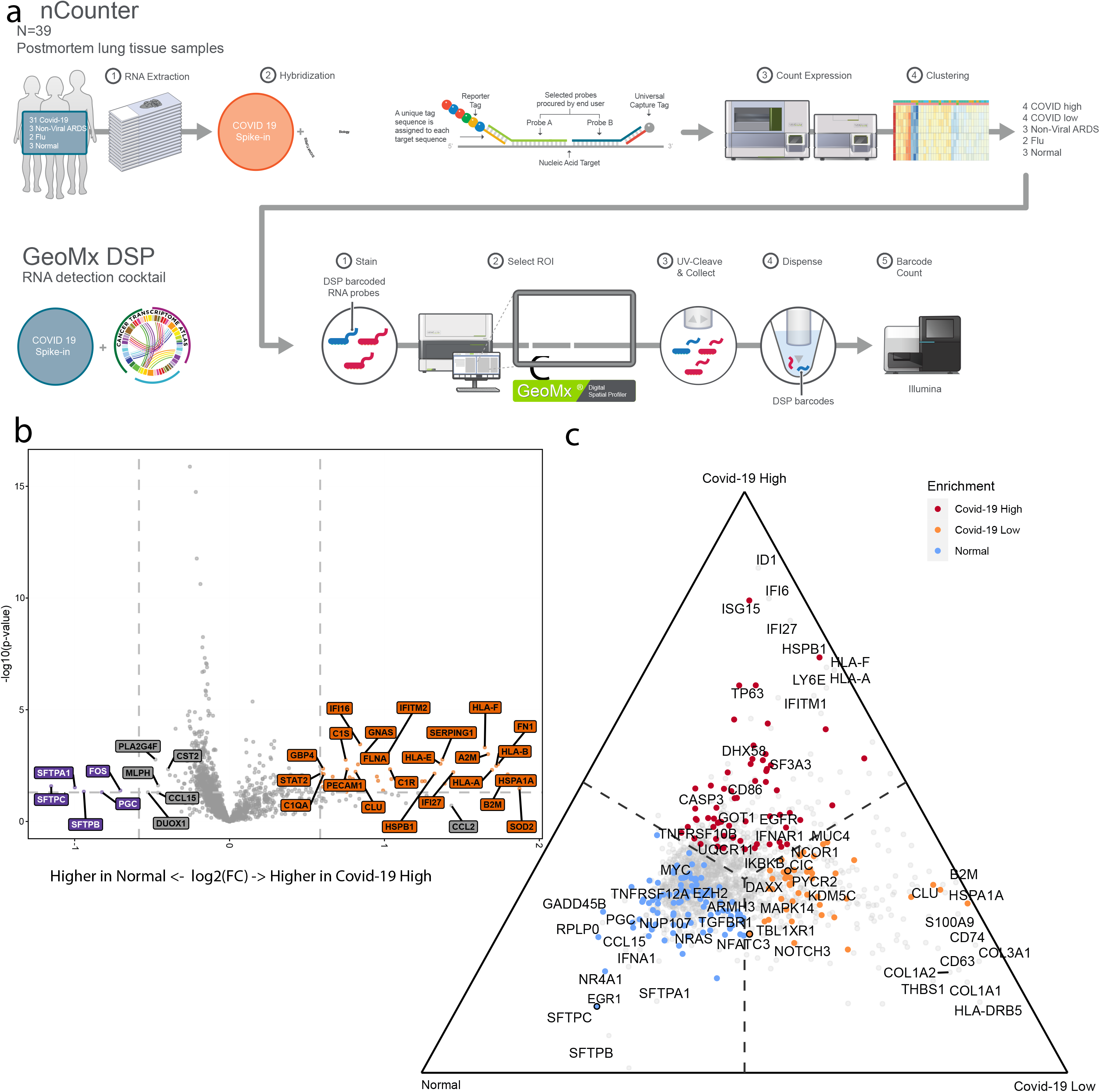
Spatial transcriptomics identifies differences between COVID-19 high and COVID-19 low relative to Normal lung. **a.** Diagram showing the two-pronged approach used to identify high and low SARS-CoV-2 viral groups from a larger cohort followed by spatially resolved transcriptomics of a subset of samples using GeoMx. **b.** Volcano plot showing differences between Normal and COVID-19 high samples after accounting for compartmental differences. Top genes, in terms of p-value or fold change (FC) are indicated in grey and COVID-19 spike-in genes are labeled in black. **c.** Ternary plot of a combined analysis of COVID-19 high, COVID-19 low, and Normal where genes are projected away from the center based on their marginal means. Genes upregulated in a single group approach that group’s corner. Top genes in terms of p-value or FC are labeled. Genes with color were significant with p-value < 0.05. Genes with outlines are significant after correcting for multiple hypothesis testing.

**Table 1.** Number of samples in this study broken down by tissue and analysis type.

|  | **GeoMx DSP**<br>**Pts (ROIs, AOIs)** | **nCounter IO360** |
| --- | --- | --- |
| COVID high | 4 (86, 86) | 8 |
| COVID low | 4 (94, 94) | 23 |
| ARDS | 3 (63, 67) | 2 |
| Flu | 2 (42, 46) | 2 |
| Control | 3 (60, 64) | 3 |
|  | **RNA-Seq** |  |
|  | **COVID+** | **COVID-** |
| Lung | 36 | 4 |
| Heart | 36 | 5 |
| Liver | 35 | 5 |
| Lymph Node | 22 | 5 |
| Kidney | 34 | 3 |
| NP Swab | 213 | 517 |

As shown in **Figure 1b**, we observed significant differences between normal and COVID-19 high samples, even after accounting for compartmental variability. These differences included a decrease in expression of SFTPA1, SFTPB, and SFTPC (markers associated with alveolar type 1 and type 2 cells), as well as an enrichment for genes associated with basal cells (TP63) and club cells (SCGB1A1), and several immune markers (e.g. HLA-B, HLA-E) (p-values all <0.05, mixed effects model) in COVID-19 high patients. A ternary plot (**Figure 1c**) of a combined analysis of COVID-19 high, COVID-19 low, and normal tissues - where transcripts are projected away from the center based on their marginal means - revealed upregulation of several genes enriched in each set of lung tissues. Enrichments included SFTPA1, SFTPB, and SFTPC (alveolar epithelial cell markers) in normal lungs, CLU (lung injury and repair) and S100A9 (enriched in activated macrophages) in COVID-19 low lungs, and TP63 (basal cell), ID1 (upregulated and a key regulator of lung injury and repair), and interferon regulated genes including IFI6, IFI27, ISG15, and LY6E in COVID-19 high lungs. Enrichment of interferon stimulated genes was only observed in COVID-19 high samples, which correspond to early stages of the infection, by timing of disease onset and histopathology (**Extended Data Figure 1**). Moreover, we observe enrichment of CASP3 and ID1, suggesting ongoing cellular injury and repair responses in COVID-19 high patients. In contrast, we find an enrichment of several markers of pulmonary fibrosis (e.g. CLU, COL1A1, COL1A2, and COL3A1) in COVID-19 low patients (which correspond to the later stages of infection), indicating that these are two distinct stages of infection^7^.

Next, we examined the spatial transcriptome data for differences between the COVID-19 high/low patients and the influenza lung samples. This analysis revealed a significant enrichment of THBS1 and NR4A1 in the influenza samples, both genes that have been shown to be engaged in response to influenza-induced lung injury (**Extended Data Figure 2a**). When lung tissues from COVID-19 high and low samples were compared to those from non-viral induced acute respiratory distress syndrome (ARDS), enrichment for S100A8 and S100A9 was observed, which is consistent with the significant enrichment of neutrophils in these samples and previously suggested to be a driver of COVID-19 pathogenesis (**Extended Data Figure 2b**)^8,9^. Importantly, an enrichment for genes involved in lung injury and repair (ID1) as well as those involved in type 1 interferon responses including IFI6, IFI27, ISG15, and LY6E in COVID-19 high lungs is observed even when compared to influenza infection of the lung (**Extended Data Figure 2a)** and non-viral lung injury (**Extended Data Figure 2b**). Similarly, we find enrichment of several markers of pulmonary fibrosis (e.g. CLU, COL1A1, COL1A2, and COL3A1) in COVID-19 low samples as compared to non-viral and viral ARDS samples, exemplifying the profound lung injury and fibrosis during later stages of COVID-19 infection.

### COVID-19 specific changes in intrapulmonary and viral heterogeneity

For each ROI, count estimates of 15 distinct cell types were imputed based on gene expression profiles from the Human Cell Atlas (HCA) adult lung dataset, including a neutrophil profile derived from snRNA-seq of lung tumors (see Methods). Consistent with other studies^10,11^, we observed that COVID-19 was associated with an increase in tissue infiltrating immune cells, including T cells, NK cells, monocytes, and macrophages (**Figure 2a**). Some immune cell types, such as monocytes, NK cells, and regulatory T cells, showed a statistically significant increase only in the COVID-19 high condition (**Figure 2**). Also, while fibroblasts and endothelial cells increased in both COVID-19 high and low samples (52% and 65% increase, respectively), type 1 and type 2 alveolar epithelial cell proportions decreased (26% and 16% decrease, respectively), reflecting the ongoing tissue remodeling or selective epithelial cell death induced by infection (**Extended Data Figure 3**). Using an orthogonal approach, we stained the lung tissues with Masson’s trichrome and observed a statistically significant increase in cellular collagen-rich areas, confirming the increase in lung fibroblasts (**Extended Data Figure 4**). As shown in the cluster analysis these cell type counts and proportions distinguish between the normal vs. COVID-19 (high and low) lungs regardless of lung structure origin (**Figure 2b**), indicating that the SARS-CoV-2 infection is altering the cellular landscape and composition of the lung tissue.

**Figure 2.**
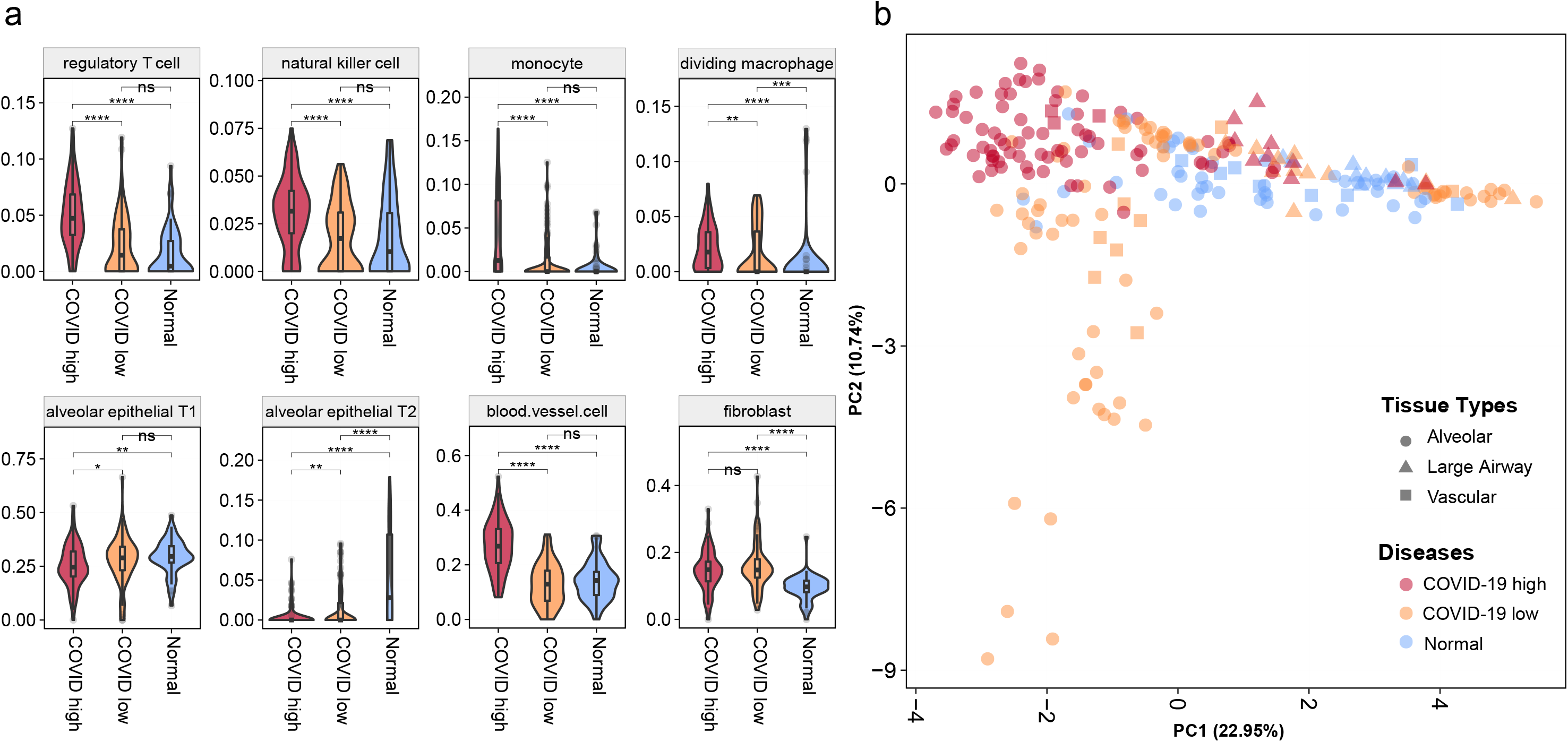
Deconvoluted cell type proportion estimates and comparison. **a.** Cell type proportions between normal vs. COVID-19. The median and quartiles are noted by the box plot inside. P-value two-tailed t-tests were done to compare the means (ns: non-significant, *:p <= 0.05, **: p <= 0.01, ***: p <= 0.001, and ****: p <= 0.0001). **b.** PCA analysis of the cell proportions. Colors denote disease conditions (normal, COVID-19 high and low), while shapes show the tissue types (alveolar, large airway, and vascular regions).

Given the observed changes in cell proportions that are induced during SARS-CoV-2 infection, we next examined the impact on the cell-to-cell interaction landscape as a metric of intrapulmonary cellular heterogeneity (**Figure 3**). Pairwise correlations of all detected cell types under five different conditions (COVID-19 high and low, flu, ARDS, and normal) were calculated and visualized (**Figure 3a**). In normal lungs, we observed three clear cellular correlation clusters (1) monocytes, fibroblasts, T and NK cells, (2) neutrophils and ciliated cells, and (3) pDCs, macrophages, and B cells. While perturbations of these cellular interactions were observed across all injury conditions, the NK-T cell correlation was lost only in the COVID-19 high patients, and not present in COVID-19 low patients, which corresponds to the later stages of infection (**Figure 3a-b**). We then quantified the correlation differences in the lungs’ cellular landscape between COVID-19 high vs. COVID-19 low patients, which indicated that the greatest changes were in the monocyte-T correlations and dendritic-neutrophil correlations (**Extended Data Figure 5a**), further supporting the view that SARS-CoV-2 specific T cell activities may be disrupted. Of note, it may be possible that the changes in correlation reflect both the generation of long-term memory as well as T cell mediated killing of infected epithelial cells^12^.

**Figure 3.**
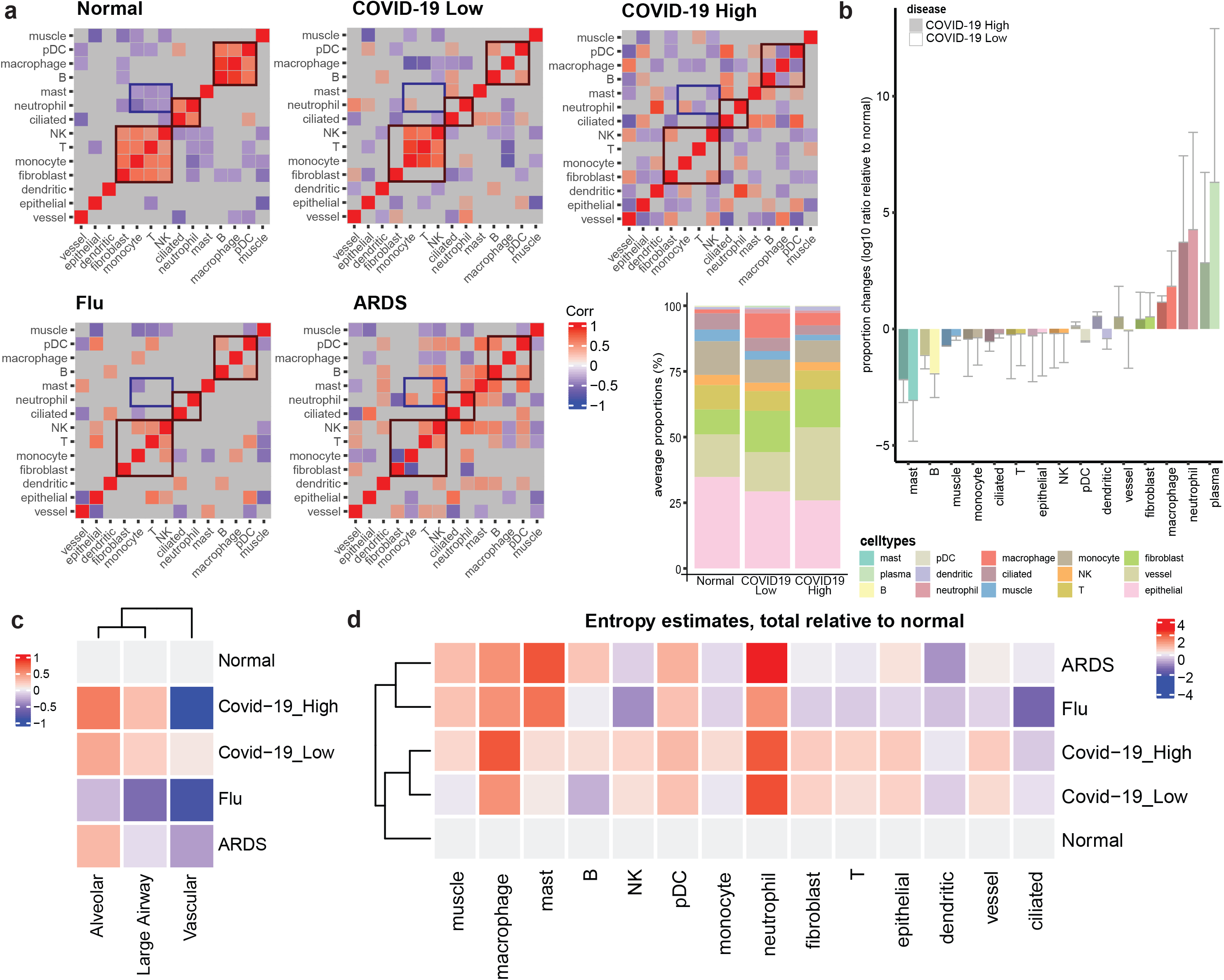
Cell Interaction Displacement in COVID-19. **a.** Correlation matrix of pairwise cell-type correlations. Statistically insignificant correlations were not displayed (grey area, p-value cutoff 0.05). **b.** average proportion changes of the cell types relative to normal. The cell types were ordered by average increase in proportions (in the plot: zoom-in of the cell types excluding neutrophils and plasma cells). Error bars indicate 0.5*SD. On the left, stacked bar plot depicts the overall proportions by conditions **c.** Entropy estimates of the tissue types, and **d.** of the cell types within the conditions

When next plotted the average proportions of each cell type across the COVID-19 disease states (COVID-19, Flu, ARDS). Macrophages and neutrophils populations were found to be much higher in the COVID-19 lungs (both high and low), while T, monocyte, and epithelial cells were much lower compared to normal. The entropy estimates of the given cell types (**Figure 3c**) showed that macrophages and neutrophils commonly displayed increased heterogeneity (across ROIs and patients) across all injury conditions. Variances of fibroblast, epithelial, vessel, and T cell populations were bigger in COVID-19 high and low lung tissues compared to flu infection and ARDS, suggesting an increase in the cellular heterogeneity. When comparing the entropies by tissue types, nearly all COVID-19 positive ROIs showed increase in heterogeneity of the cell populations, with the sole exception being vascular regions with high COVID-19 (**Figure 3d**). The decrease in entropy in the vascular regions of COVID-19 high condition is mainly from the decrease of fibroblasts, epithelial, T, and NK cells (**Extended Data Figure 5b**). The signals from B cells are also specific to large airway tissues, and this observation is consistent with the cell fraction increase in large airway ROIs of COVID-19 samples (**Extended Data Figure 5b**). Single-sample Gene Set Enrichment Analysis (ssGSEA) of macrophage-, neutrophil-, and T cell regulatory pathways showed enrichment in COVID-19, even when compared with flu and ARDS, including macrophage activation and apoptotic process (1.302 and 2.809 fold increase in averaged ssGSEA scores relative to normal, with p-values of 2.91e-08 and 0.001) in COVID-19 high. (**Extended Data Figure 6a-c)**.

SARS-CoV-2 RNA reads were aligned to the SARS-CoV-2 genome and the number of reads was discerned across multiple tissues including the lung, lymph node, kidney, liver, and heart (**Extended Data Figure 7a-b**) but mostly in the lung. Viral reads were robustly detected in nasopharyngeal samples from COVID-19 patients, consistent with the published reports^13^. Normalized coverage values in SARS-CoV-2 positive autopsy tissue samples revealed detection bias towards the SARS-CoV-2 3’ end consistent with the known life cycle of the virus (**Extended Data Figure 7a-b)** ^13,14^. Reconstruction of the viral genomes revealed known and unknown variants common to many patients (**Extended Data Figure 7c),** and some evidence of intra-host variability.

### Multi-organ analysis from patient autopsy samples

We next used shotgun metatranscriptomics (total RNA-seq) for host and viral profiling on 39 patients that died from COVID-19, including 190 organ-specific tissue samples from the respective autopsies and healthy controls from organ donor remnant tissues (n=3). We examined the COVID-19 specific host responses and transcriptome changes across various organs (heart, kidney, liver, lung, and lymph nodes) to ascertain the differentially expressed genes (DEGs) between COVID-19 high, COVID-19 low, and control sample sets (q-value <0.01 expression fold-change >1.5-fold, DESeq2, **Figure 4a-d**). Pathway enrichment analysis revealed significant changes (q-values <0.01) in pathways for viral infection (Regulation of Viral Genome Replication and Viral Entry into Host Cell), immune response (Regulation of Type 1 Interferon Response and Regulation of Tyrosine Phosphorylation of Stat Protein, GO_Regulation of Toll Like Receptor Signaling Pathway, **Supplementary Table 1**).

**Figure 4.**
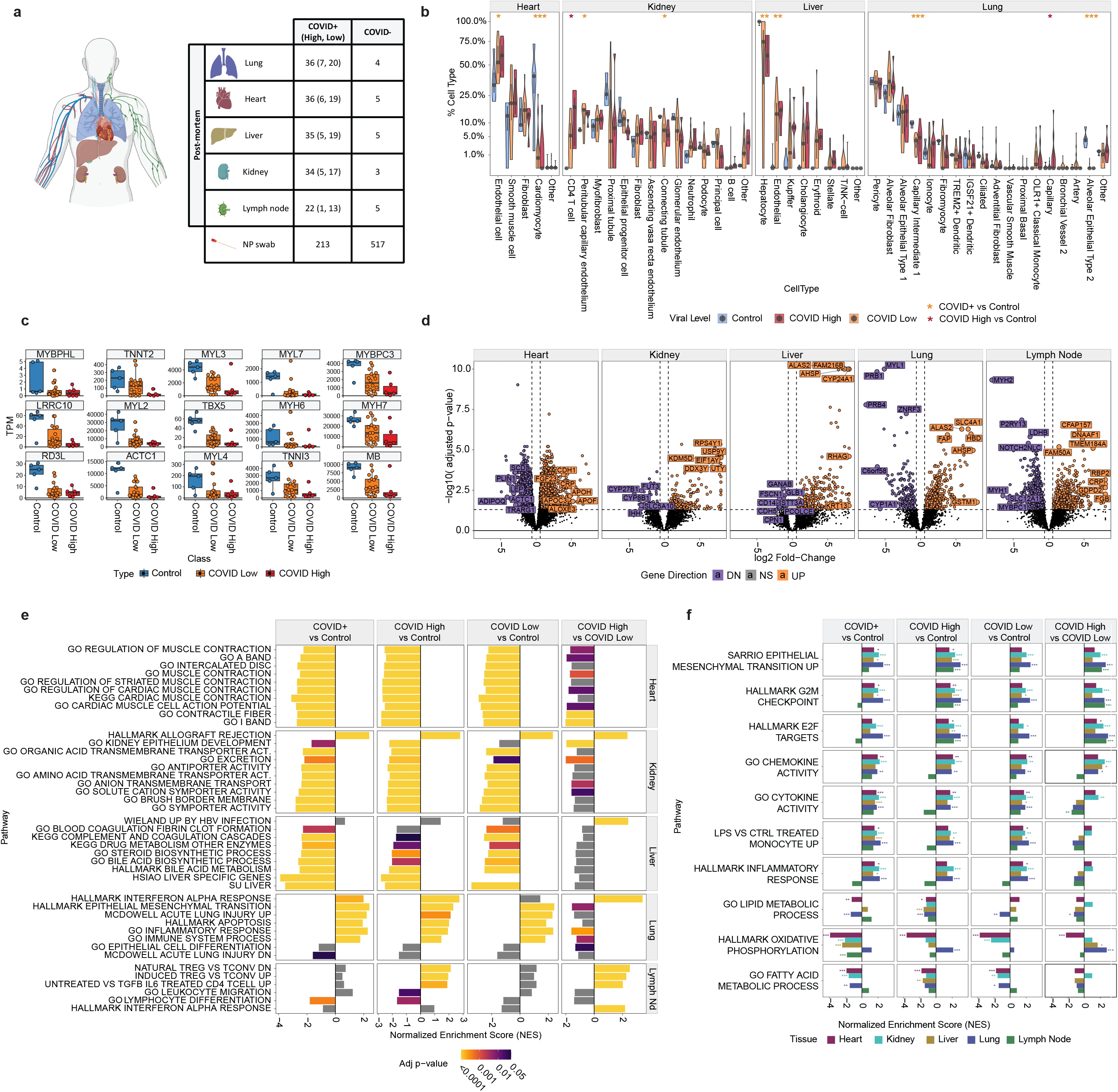
Tissue- and duration-specific dysregulation of gene expression from SARS-CoV-2 infection. **a.** Sample overview by tissue for RNA-seq experiments. **b.** Cellular deconvolution distribution violin plots for the heart, kidney, liver, and lung. COVID-19 high (red), COVID-19 low (orange), normal tissues (blue) are shown in square root scale, as well as **c.** specific gene expression distributions for cardiomyocyte-related genes. **d.** Volcano plots of the COVID-19 positive vs. normal tissues are shown for the five tissues from autopsy: heart, kidney, liver, lung, lymph node. Differentially expressed genes (>1.5-fold, q-values <0.05, DESeq2) are shown as purple spots (down-regulation) and orange (up-regulation). **e.** Select pathways that show significant differences in different tissues are shown, with the statistical significance shown as a color range (legend, GSEA permutation test and adaptive multilevel splitting Monte Carlo method), and non-significant differences shown in grey. **f.** The pathways that show significant differences in all or majority of the five tissues are shown, with statistical significance from GSEA testing across five tissues (color in legend) and x-axis for the normalized enrichment score for COVID-19 positive vs. Control, COVID-19 high or low vs. Control, and COVID-19 high vs. COVID-19 low comparisons.

We observed that each tissue showed its own distinct transcriptional disruption due to the infection, with the lymph node exhibiting the greatest number of DEGS when compared to controls (in both COVID-19 high and COVID-19 low patients). Of note, both tissue-specific and pan-tissue disruptions of normal expression programs were observed (**Figure 4a-d**), and these were then summarized using GSEA (**Figure 4e-f**). Some pathways were consistently dysregulated in all tissues during early infection (COVID-19 high), such as G2M checkpoint (q-values of 6.4 × 10^−19^, 3.0 × 10^−5^, 1.2 × 10^−8^, 9.7 × 10^−8^, and 0.002 for lung, liver, kidney, lymph node, and heart respectively, **Supplementary Table 1**), E2F targeting (q-values of 2.8 × 10^−26^, 0.00672, 1.9 × 10^−7^, 9.9 × 10^−9^, 0.047, respectively), and epithelial mesenchymal transition (EMT) (q-values of 2.1 × 10^−22^, 3.0 × 10^−4^, 2.9 × 10^−7^, 1.4 × 10^−7^, 0.0287, respectively), but late infection (COVID-19 low) gene networks showed more inter-tissue heterogeneity in their disrupted pathways, including cytokine activity and inflammatory response. However, in both the COVID-19 high vs. low comparisons, the G2M checkpoint and E2F networks were consistently upregulated, indicating a core, persistent set of dysregulated cell cycle regulation genes during early and late stages of infection.

The DEGs and GSEA results were then examined for the largest differences between infection level and stage (COVID-19 high, early infection vs. COVID-19 low, late infection). Interestingly, the heart tissues showed the largest transcriptional differences, revealing that the later stage of the infection had a much greater impact on cardiac tissues (**Figure 4**). To place these results into greater context, we compared DEGs from each tissue to RNA-seq data from nasopharyngeal (NP) swabs, previously described in Butler et al^4^, as well as RNA-seq data from a publicly available dataset on monocytes from COVID-19 positive and negative patients (**Extended Data Figure 10**). The majority of the tissues with high viral load were significantly positively correlated with the DEGs in the NP swab samples (q-value <0.01), compared to normal/negative patients. However, in contrast, the later infection (low viral load) patients’ tissues showed negative correlation with the NP swab samples, indicating that the systemic impact of SARS-CoV-2 can be missed when not considering the biological impact on different organs.

Despite significant transcriptional differences in the lung, there were no clear gross pathologic differences in bulk tissue or cell surface proteins (**Extended Data Figure 8a**, **8b)**. Thus, to create a more fine-grained analysis of the cellular gene expression state in each tissue, we used the cell deconvolution algorithm MuSiC on each tissue’s RNA-seq data. The MuSiC results showed distinct disruptions of the transcriptional programs for each tissue in the COVID-19 patients and gain or loss of cell types (**Figure 4b**). Specifically, the kidney and liver showed a loss of proximal tubule in the kidney and hepatocyte marker expressions in the liver (**Extended Data Figure 4a, 10**), but an increase in T cells found in the kidney and liver. The lung showed a loss of the capillary intermediate cells and alveolar epithelial cells. Finally, the heart showed a striking, near-complete loss of the cell signatures for cardiomyocytes (**Figure 4c**), in both COVID-19 high and low samples (**Extended Data Figure 4a, 9**), despite no obvious gross or histologic changes in the heart (**Extended Data Figure 8b**).

Given the changes in expression phenotypes of the infected tissues, we next examined the splicing differences induced in each tissue relative to controls. Quantified transcript counts from RNA-seq were used to estimate isoform differences with Isoform Switch AnalyzeR (ISAR)^18^. Each tissue was examined for transcript switches across nine isoform disruption categories, such as loss of function, disrupted open reading frames, and nonsense mediated decay (FDR <0.05, **Figure 5a**). Splicing alterations were most pronounced in the heart and lung tissues (7/9 isoform types significantly altered), but less significantly in the lymph nodes (2/9) and liver (0/9) again highlighting a particularly disruptive set of transcriptional changes in the heart and lung. Next, alternative splicing was analyzed within each tissue (**Figure 5a**) from the ISAR data, showing significant exon skipping in the lung and increased exon skipping events in the heart, and more alternate start and stop sites within the heart, lung, and lymph nodes (FDR<0.05). No significant isoform switching or alternative splicing was detected in the kidney, despite significant disruption in expression profiles.

**Figure 5.**
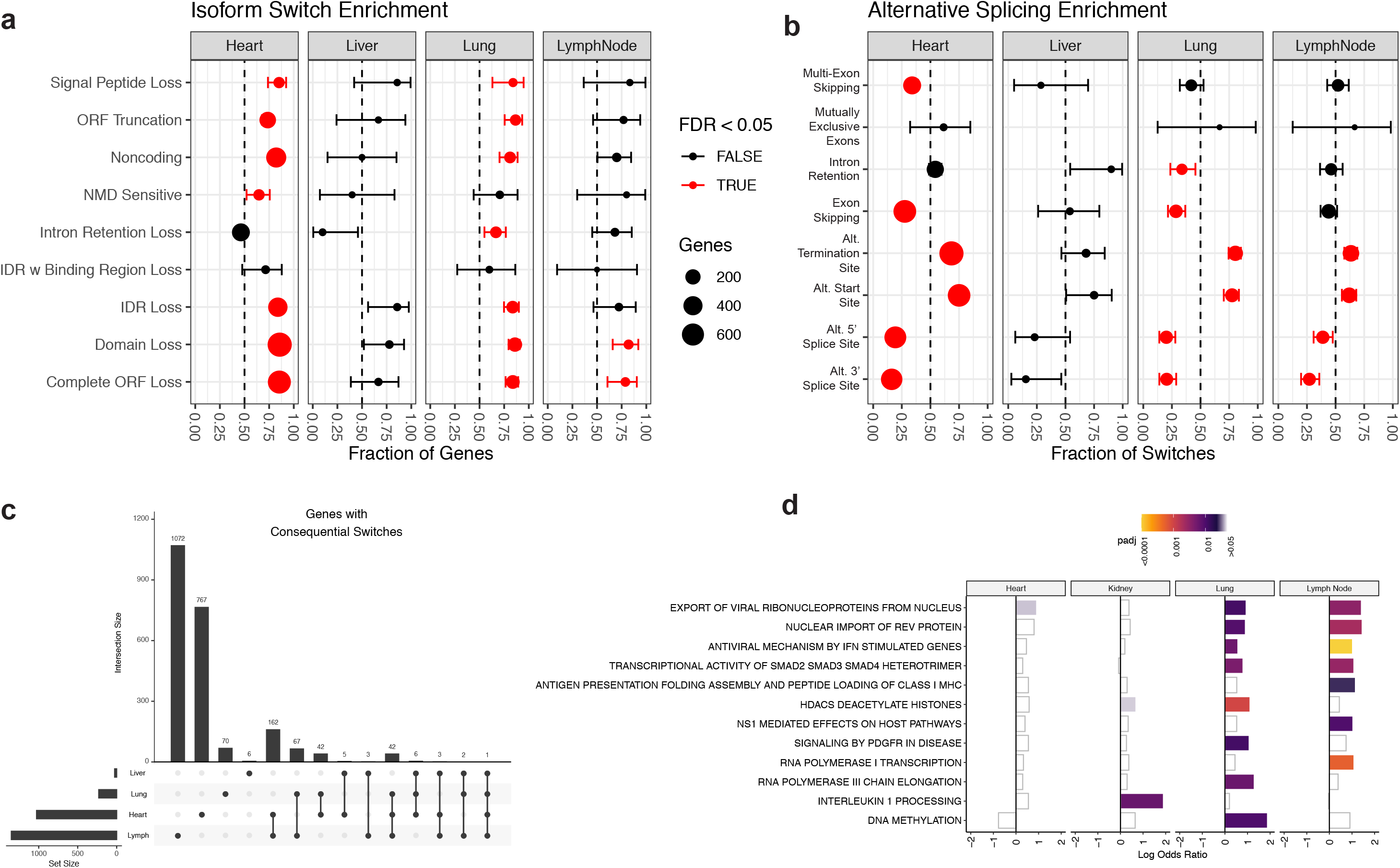
Tissue-specific splicing dysregulation from SARS-CoV-2 infection. **a.** Enrichment of isoform switching types across tissues. ORF=Open Reading Frame; NMD=Nonsense-Mediated Decay; IDR=Intrinsically Disordered Regions. **b.** Enrichment of alternative splicing across tissues. **c.** UpSet plot showing counts of genes with consequential isoform switching per tissue. **d.** Log odds ratio of select significantly dysregulated pathways across tissues, colored by adjusted p-value.

To put these differences into broader context, we then examined the intersection of specific splicing-disrupted genes across tissues and pathways (**Figure 5c,d**). The greatest number of genes with consequential isoform switches (significant genes from **Figure 5a,b**) was found in the lymph node (1353) and heart (1029), followed by lung (231) and liver (27). Most splice-disrupted genes were unique to a given tissue, but the greatest overlap was observed between the heart and the lymph node (162 genes). Next, pathway enrichment was calculated with PathwaySplice^19^, using canonical pathways from the Reactome database (**Figure 5d**). In the tissues with significant isoform switches, the greatest number of significantly disrupted pathways (adjusted p-value < 0.05) occurred in the heart (97), followed by the lymph node (72) and lung (65), indicating a stronger pathway disruption effect in the heart than the lymph node and lung. Of note, the top dysregulated pathways in these tissues were related to viral responses, including the export of viral ribonucleoproteins and antiviral mechanism by interferon-stimulated genes, reflecting a broad, but tissue-specific, transcriptional response to viral infection (**Extended Data Figure 10**).

### Characterization of SARS-CoV-2 related TCEM repertoires and abundances

Finally, to examine the impact that COVID-19 has on T cell receptors in the patients that could be found in each tissue, we assembled CDR3 sequences from the patient-specific RNA-Seq data and derived a set of T Cell Expressed Motifs (TCEMs); TCEMs are amino acid sequences that can be recognized as an antigen by a T Cell when presented by an MHC. We mapped these TCEMs to the SARS-CoV-2 reference genome, to quantify the likely response of the T cells to the virus (**Figure 6)**. When compared the abundance of TCEMs that mapped to the SARS-CoV-2 genome, the abundance of TCEMs that mapped to SARS-CoV-2 was significantly higher in cases than in controls (3.7x increase using one-sided T-test, p-value of 1.24e-12, **Figure 6a** and **Supplementary Table 3**).

**Figure 6.**
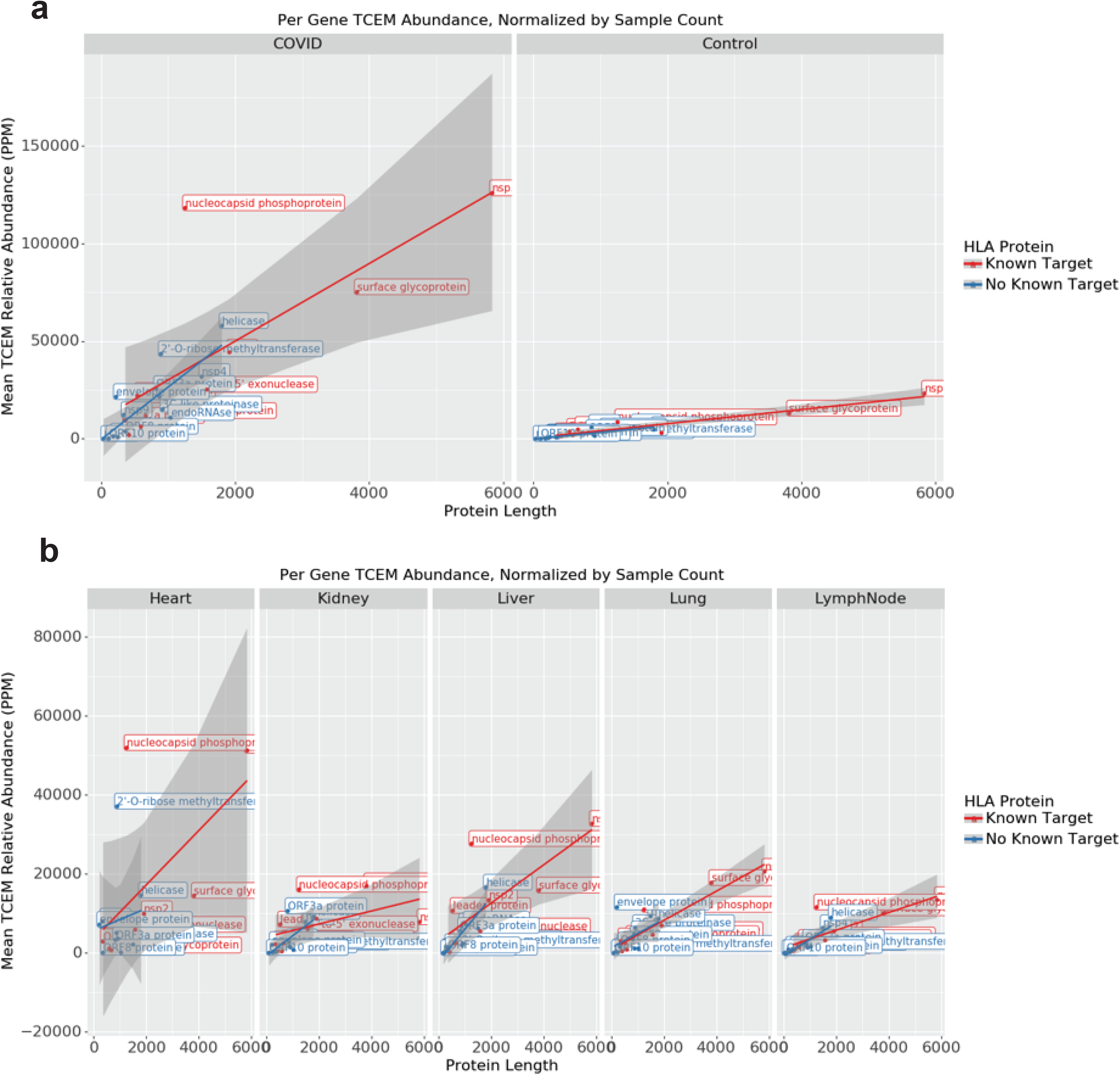
TCEMs match sites throughout the SARS-CoV-2 genome. **a.** abundance of TCEMs that mapped to the SARS-CoV-2 genome in both our case and control samples. **b.** Average abundance of TCEMs across different tissue types. Coefficients of fit and p-values for each line in both panels are listed in **Supplementary Table 3**, and confidence intervals are visualized in grey area.

We next considered variable (non-uniform) TCEM abundance across different tissue types. By taking the average abundance of TCEMs that mapped to each gene in the SARS-CoV-2 genome across all patients for each tissue type (lung, liver, heart, kidney, and lymph nodes, **Figure 6b**), we found that TCEMs matching SARS-CoV-2 are differentially abundant by tissue types. The majority (2592/3174, 81.7%) of unique TCEMs were found in samples from lymph nodes, consistent with their importance in immune function. Despite the largest number of unique TCEMs, those from the lymph nodes were typically low abundance compared to other tissues, which may be evidence of T Cell maturation before effective clones are found and distributed across the body. By contrast TCEMs from heart samples were highly abundant and less diverse, indicating a possible targeted response, which also provides further evidence of the distinct cardiovascular COVID-19 phenotype of heart damage.

## Discussion

In this study, we established a clinical analytical pipeline to collect and examine autopsy samples to elucidate and compare the spatial transcriptional landscape induced by SARS-CoV-2, influenza, and ARDS. First, we found that the lung and other tissue samples demarcated COVID-19 patients into two groups, based on the amount of SARS-CoV-2 RNA present: COVID-19 high versus COVID-19 low. Lung samples examined using two techniques (Nanostring nCounter or GeoMx DSP) both support the correlation that high SARS-CoV-2 viral loads correlate with early COVID-19 disease while low SARS-CoV-2 viral loads correlate with late COVID-19 disease. The nature of the COVID-19 disease and the impact on the lung (and other tissues) appears to be defined principally by the presence of SARS-CoV-2 viral RNA.

Patient lung tissue samples containing significant levels of SARS-CoV-2 showed enrichment for genes related to a variety of immune markers specific to certain immune cells and lung injuries, as well as enriched for interferon stimulated genes (e.g. *IFI27*, *IFITM1*, and *LY6E*) and macrophage activation (*S100A9*, *TYMP*, and *SERPING1*). In contrast, patient lung tissue samples containing low levels of SARS-CoV-2 RNA show enrichment for *COL1A1* and other markers of pulmonary fibrosis. Compared to other viral related diseases (influenza), COVID-19 tissue samples still show enrichment for genes involved in lung injury and repair and interferon signaling genes, but the COVID-19 tissue samples also show enrichment of several markers of pulmonary fibrosis. Of note, COVID-19 (high), flu, and ARDS each show differential *HLA-B* and *-C* expression, which are known mediators of natural killer and T cell activation^20,21^ and which can mediate host risk of infection^4^. Enrichment for *HLA-DRB5,* whose expression and specific gene polymorphisms are associated with pulmonary fibrosis and severity^22,23^. Comparing across different disease types, all diseases—COVID-19 (low), influenza, and ARDS showed enrichment for *DMBT1*, a gene known to be upregulated and dysregulated in pulmonary injury and fibrosis^24,25^. Virus-related diseases (COVID-19 high and flu) particularly showed significant change of expression in lung epithelial cell-related transcripts (i.e. *ACTB*, *C1R*, and *FN1*), and such changes are known markers of the lung injury gene signature^26^.

The spatial analysis platform (GeoMx) enabled a novel analysis of the impact of the disease across entire tissues. Consistent with recent reports from bulk cellular profiling, we observed an increase in the immune cell types and fibroblasts in COVID-19, high but a decrease in alveolar epithelial cells^27^. In COVID-19-low condition, the proportions of some immune cells (i.e. monocytes, NK cells, or regulatory T cells) were similar to normal, but the fibroblasts and vessel cells still exhibited an increase in COVID-19. Some of these cell types form a “cellular correlation cluster,” and these clusters of cell-cell interactions are uniquely disrupted in COVID-19 (relative to flu and ARDS), particularly in COVID-19 high. While macrophages and neutrophils showed an increase in entropy across all lung-related injury conditions, NK and T cells only showed an increase in COVID-19 conditions. While few studies have interrogated the tissue environments; multiple studies have examined the changes occurring during COVID-19 infection in the peripheral blood and have identified poor T cell responses and T cell dysregulation^28–31^. Together, these findings highlight the robust and dynamic nature of SARS-CoV-2 engagement with tissue homeostatic processes and that the stage of COVID-19 infection impacts the pathophysiological landscape of the lung.

When these spatial omics data were compared to the multi-organ bulk RNA-seq data from the autopsy issues, confirmatory as well as additional signatures of COVID-19 disease were found. First, the fibroblasts and immune cell types such as macrophages increased in most tissues, while alveolar epithelial types 1 and 2 cells in the lung showed a decrease in COVID-19 (relative to controls). Interestingly, other organ types such as heart or kidney also showed similar trends in COVID-19. Increase in fibroblasts, endothelial, and immune cells may be impacted by a variety of immune activations within each organ, particularly as s long-term response to the infection. This observation may also be related to the decrease in the characteristic transcriptomic signatures of the main cell type within each organ, which may contribute to the morbidity and mortality of COVID-19. For example, the reduction in cardiomyocyte cell fraction and disrupted splicing isoforms within heart tissues was concomitant with a reduction in several transcripts encoding sarcomeric and contractile proteins, regardless of viral level^32,33^, representing a persistent transcriptional perturbation and cardiac-specific impact of COVID-19.

These data support the view of distinct biological responses to the stages of SARS-CoV-2 infection, and this is buttressed by orthogonal data. For example, DEGs observed from NP swabs showed high correlation to tissue specific DEGs in the early stages of infection, but very little correlation later infection. To explain this phenotype, recent evidence has suggested that monocytes can migrate into tissues as the SARS-CoV-2 infection progresses^15^, and this response can potentially explain the different correlations. It has also been reported that monocyte depletion/migration is associated with kidney disease, inducing lupus like symptoms^16^, which could potentially explain the correlation with kidney tissue. For lymph nodes, there exists evidence in the literature that SARS-CoV-2 will impact the lymph nodes at an early stage of infection potentially causing T cell lymphopenia and possibly responsible for focal necrosis seen in the lymph nodes^17^. Nonetheless, the lung, heart, and lymph nodes were the tissues most disrupted by infection.

In addition to our observations, there are several papers indicating the connections of cytokine storms with macrophage activation syndromes and/or other imbalance of the immune system environment^34,35^, which also can be variable across tissues. To explore these connections, we examined the relative abundance in TCEMs mapped to the SARS-CoV-2 genome in our RNA-seq results. Organ-wide TCEM mapping also revealed a wide variety of TCEM sites in heart, liver, or lung and showed high similarity within the body sites or patients. Indeed, the ssGSEA from the lung specific dataset of COVID-19 on the T cell mediated pathways revealed distinct from that of ARDS, flu, and the normal samples (**Extended Data Figure 6c**). The expression signature was particularly prominent for T cell proliferation, chemotaxis, and activation, and this impact was most evident in the lymph node and the heart.

Overall, these data represent one of the largest autopsy series of COVID-19 and synthesizes several orthogonal methods, including: bulk transcriptomics, digital spatial transcriptomics, multiple imaging technologies, and novel computational analysis (TCEMs, ISAR) to build a map of SARS-CoV-2 pathophysiology. The vast majority of COVID-19 data previously published has been from NP swab or peripheral blood, and here we provide evidence that these data cannot capture the key, tissue-specific impacts of the disease, especially in late infection. Moreover, given the ability to also use these data to map the distinct viral variants as the pathogen moves through the body, our data also provide additional support of intra-host viral diversity for SARS-CoV-2, including variations specific to distinct tissues^36^, which can help guide future studies and treatments. This molecular map of COVID-19 serves as an atlas for the community and can inform future studies into COVID-19 progression and SARS-CoV-2 pathology.

## Acknowledgements

We thank the patients, their families, and healthcare workers fighting the COVID-19 pandemic. This work was supported by the NCI (R01CA234614) and NIAID (2R01AI107301) and NIDDK (R01DK121072 and 1RO3DK117252) to Department of Medicine, Weill Cornell Medicine (R.E.S.). R.E.S. is supported as an Irma Hirschl Trust Research Award Scholar. A.B. is supported by supplemental funds for COVID-19 research from Translational Research Institute for Space Health (TRISH) through NASA Cooperative Agreement NNX16AO69A (T-0404) and also further funding was provided by KBR, Inc. Sequencing of some samples was performed at the New York Genome Center (NYGC) as part of the COVID-19 Genomic Research Network (CGRN) with funds generously provided by NYGC donors. We would like to thank the Epigenomics Core Facility at Weill Cornell Medicine, the Scientific Computing Unit (SCU), as well as the Starr Cancer Consortium: I9-A9-071, I13-0052, the NIH: R21AI129851, R01MH117406, R01CA249054, R01AI151059, P01CA214274, the Leukemia and Lymphoma Society (LLS) MCL7001-18, LLS 9238-16, LLS-MCL7001-18).

## Author Contributions

RES, AS, and CEM conceived and designed the experiments. JP, TH, SW, YK, JRe performed GeoMx analysis and figures. CMe and JF performed the RNA-sequencing bioinformatics analyses and statistical investigations, with contributions from DD. ACB was involved in autopsy tissue procurement, pathology evaluation, GeoMX ROI selection and trichrome quantitation. RES processed and analyzed samples and clinical data, with contributions from YB, VC. DJB, CMoz, EA, MM, SL, MS, AMM, IH, SW, AC, PV, MS, ML, LFW, MC, HR, NT, organized and worked with nasopharyngeal swabs and data. HG, SF, AC, MCZ, SG, AB, DT, ASB, US, ESW, JS performed the RNA-sequencing and worked on data analysis, interpretation, figures, with contributions from EA, MI, AS, JRo, SSa, SSh, AJK, OE. All authors discussed the results and contributed to the final manuscript.

## IRB Statement

Tissue samples were provided by the Weill Cornell Medicine Department of Pathology. The Tissue Procurement Facility operates under Institutional Review Board (IRB) approved protocol and follows guidelines set by Health Insurance Portability and Accountability Act (HIPAA). Experiments using samples from human subjects were conducted in accordance with local regulations and with the approval of the IRB at the Weill Cornell Medicine. The autopsy samples are considered human tissue research and were collected under IRB protocols 20-04021814 and 19-11021069. All autopsies have consent for research use from next of kin, and these studies were determined as exempt by IRB at Weill Cornell Medicine under those protocol numbers.

## Data Availability

All the raw sequence files and metadata for specimens, including per-run metrics and QC data, have been submitted to the database of Genotypes and Phenotypes dbGAP (accession #38851 and ID phs002258.v1.p1). All relevant data are available at these databases: https://www.ncbi.nlm.nih.gov/projects/gap/cgi-bin/study.cgi?study_id=phs002258.v1.p1. Nanostring GeoMx data are also deposited in the GEO database). Processed bulk RNA-seq data is also available online (https://covidgenes.weill.cornell.edu/).

## Methods

### Patient sample collection

All autopsies are performed with consent of next of kin and permission for retention and research use of tissue. Autopsies were performed in a negative pressure room with protective equipment including N-95 masks; brain and bone were not obtained for safety reasons. All fresh tissues were procured prior to fixation and directly into Trizol for downstream RNA extraction. Tissues were collected from lung, liver, lymph nodes, kidney, and the heart as consent permitted. For GeoMx, RNAscope, trichrome and histology tissue sections were fixed in 10% neutral buffered formalin for 48 hours before processing and sectioning. These cases had a post-mortem interval of less than 48 hours. For bulk RNA-seq tissues, post-mortem intervals ranged from less than 24 hours to 72 hours (with 2 exceptions - one at 4 and one at 7 days - but passing RNA quality metrics) with an average of 2.5 days. All deceased patient remains were refrigerated at 4C prior to autopsy performance.

### Spatial Transcriptomics Analysis

Gene Expression profiling of freshly extracted RNA from formalin fixed paraffin-embedded (FFPE) lung samples was performed using the NanoString PanCancer IO360 panel with custom probes for SARS-CoV-2 viral genes. After normalization, high and low COVID-19 clusters were identified by unsupervised analysis, and samples from each cluster were selected for additional profiling. GeoMx Digital Spatial Profiling (DSP) was performed on these samples, and control samples from non-viral ARDS, Flu, and normal lung tissues following standard protocols using the COVID-19 Immune Response Atlas^37^. Samples were stained with immunofluorescent antibodies for CD68, CD45, PanCK, and DNA (Syto-13). Regions profiled included vascular zone, large airway, alveoli zone, and IF-guided segments focused specifically on macrophages. Samples were sequenced on an Illumina NextSeq, processed and filtered for quality as described in supplementary methods. Differential expression was assessed on the resulting normalized data using mixed effect models, accounting for intra-patient heterogeneity to assess differences between COVID-19 high and low populations, and among distinct tissue structures profiled. Cell deconvolution of the GeoMx data was performed using the SpatialDecon R package^38^. Gene set enrichment analysis (GSEA)^39^ was performed to qualify coordinate gene expression changes quantified during differential expression analysis.

### Pairwise correlations of cell types by conditions

Correlation matrix visualizes the Pearson correlation coefficient by cell types within each disease condition. Statistically insignificant correlations (p-value bigger than 0.05) are filtered and identified clusters of positive and negative correlation is marked. The correlations from COVID-19 high and low are compared with normal, using R package cocor (v1.1-3)^40^. Briefly, the correlation coefficients are tested using Fisher’s r-to-Z transformation to quantify the differences between the two correlations. To quantify correlations, each data point (or correlation coefficient) corresponds to a fisher-tested correlation (z statistics and –log(P-value) for x and y axes, respectively). The entropy calculations were done with the synRNASeqNet R package (v1.0, entropyML function, https://github.com/cran/synRNASeqNet). The deconvoluted cell counts were used as an input to run maximum likelihood entropy calculations.

### GeoMx Analysis

Cell Bender was used to remove ambient RNA and other technical artifacts from the count matrices. Following CellBender, individual samples were processed using Cumulus, including filtering out cells/nuclei with fewer than 400 UMI, 200 genes, or greater than 20% of UMls mapped to mitochondrial genes.

### Antibodies

Immune Cell Profiling Panel (Core); Nanostring Technologies, Inc.; GMX-PROCONCT-HICP-12, Item 121300101, Lot# 0474026

10 Drug Target Panel; GMX-PROMODNCT-HIODT-12, Item 121300102, Lot# 0474029

Immune Activation Status Panel; Nanostring Technologies, Inc.; GMX-PROMODNCT-HIAS-12, Item 121300103, Lot# 0474032

Immune Cell Typing Panel; Nanostring Technologies, Inc.; GMX-PROMODNCT-HICT-12, Item 121300104, Lot# 0474035

Cell Death Panel; Nanostring Technologies, Inc.; GMX-PROMOD-NCTHCD-12, Lot# 0474050 MAPK Signaling Panel; Nanostring Technologies, Inc.; GMX-PROMOD-NCTHMAPK-12, Lot# 0474047

Pl3K/AKT Signaling Panel; Nanostring Technologies, Inc.; GMX-PROMOD-NCTHPl3K-12, Lot# 0474053

COVID-19 GeoMx-formatted Antibody Panel including (TMPRSS2, clone EPR3861; ACE2, clone EPR4436; Cathepsin L/V/K/H, clone EPR8011; DDX5, clone EPR7239; and SARS-CoV-2 spike glycoprotein, polyclonal); Abeam; ab273594, Lot# GR3347471-1

GeoMx Solid Tumor TME Morphology Kit; Nanostring Technologies, Inc.; GMX-PRO-MORPH-HST-12; Item 121300310

Alexa Fluor^®^ 647 alpha-Smooth Muscle Actin Antibody, clone 1A4; Novus Bio; IC1420R

Nanostring morphological and staining panels are pre-validated by the manufacturer: https://www.nanostring.com/download_file/view/2872/8714

Morphological markers were previously demonstrated in human tissue in https://doi.org/10.1101/2020.08.25.267336

### qRT-PCR

Total RNA was extracted in TRIzol (Invitrogen) according to the manufacturer’s instructions. To quantify viral replication, measured by the expression of sgRNA transcription of the viral N gene, one-step quantitative real-time PCR was performed using SuperScript III Platinum SYBR Green One-Step qRT-PCR Kit (Invitrogen) with primers specific for the TRS-L and TRS-B sites for the N gene as well as ACTB as an internal reference. Quantitative real-time PCR reactions were performed on an Applied Biosystems QuantStudio 6 Flex Real-Time PCR Instrument (ABI). Delta-delta-cycle threshold (ΔΔCT) was determined relative to ACTB levels and normalized to mock infected samples. Error bars indicate the standard deviation of the mean from three biological replicates. The sequences of primers/probes are provided in **Supplementary Table 2**.

### RNA-seq Analysis

Patient specimens were processed as described in Butler et al., 2020^4^. Briefly, nasopharyngeal (NP) swabs were collected N using the BD Universal Viral Transport Media system (Becton, Dickinson and Company, Franklin Lakes, NJ) from symptomatic patients. Total Nucleic Acid (TNA) was extracted from using automated nucleic acid extraction on the QIAsymphony and the DSP Virus/Pathogen Mini Kit (Qiagen). Autopsy tissues were collected from lung, liver, lymph nodes, kidney, and the heart and were placed directly into Trizol, homogenized and then snap frozen in liquid nitrogen. At least after 24 hours these tissue samples were then processed via standard protocols to isolate RNA.

For RNA library preparation, all samples’ TNA were treated with DNAse 1(Zymo Research, Catalog # E1010). Post-DNAse digested samples were then put into the NEBNext rRNA depletion v2 (Human/Mouse/Rat), Ultra II Directional RNA (10ng), and Unique Dual Index Primer Pairs were used following the vendor protocols from New England Biolabs. Completed libraries were quantified by Qubit and run on a Bioanalyzer for size determination. Libraries were pooled and sent to the WCM Genomics Core or HudsonAlpha for final quantification by Qubit fluorometer (ThermoFisher Scientific), TapeStation 2200 (Agilent), and qRT-PCR using the Kapa Biosystems Illumina library quantification kit.

NYGC RNA sequencing libraries were prepared using the KAPA Hyper Library Preparation Kit + RiboErase, HMR (Roche) in accordance with manufacturer’s recommendations. Briefly, 50-200ng of Total RNA were used for ribosomal depletion and fragmentation. Depleted RNA underwent first and second strand cDNA synthesis followed by adenylation, and ligation of unique dual indexed adapters. Libraries were amplified using 12 cycles of PCR and cleaned-up by magnetic bead purification. Final libraries were quantified using fluorescent-based assays including PicoGreen (Life Technologies) or Qubit Fluorometer (invitrogen) and Fragment Analyzer (Advanced Analytics) and sequenced on a NovaSeq 6000 sequencer (v1 chemistry) with 2×150bp targeting 60M reads per sample.

### Differential Gene Analysis

RNAseq data was processed through the nf-core/rnaseq pipeline^41^. This workflow involved quality control of the reads with FastQC^42^, adapter trimming using Trim Galore! (https://github.com/FelixKrueger/TrimGalore), read alignment with STAR^43^, gene quantification with Salmon^44^, duplicate read marking with Picard MarkDuplicates (https://github.com/broadinstitute/picard), and transcript quantification with StringTie^45^. Other quality control measures included RSeQC, Qualimap, and dupRadar. Alignment was performed using the GRCh38 build native to nf-core and annotation was performed using Gencode Human Release 33 (GRCH38.p13). FeatureCounts reads were normalized using variance-stabilizing transform (vst) in DESeq2 package in R for visualization purposes in log-scale^46^. Cell deconvolution was performed using MuSiC on single cell reference datasets for lung, liver, kidney, and heart^47–51^. Immune cell deconvolution was performed on lymph node samples using quanTIseq^52^. Differential expression of genes was calculated by DESeq2. Differential expression comparisons were done as either COVID+ cases versus COVID- controls for each tissue specifically, correcting for sequencing batches with a covariate where applicable, or pairwise comparison of viral levels from the lung as determined by nCounter data. In the volcano plot protein coding genes were plotted using Gencode classifications using - log10(adjusted p-value) and log2 fold-change metrics. Genes with BH-adjusted p-value < 0.01 and absolute log2 fold-change greater than 0.58 (at least 50% change in either direction) were taken as significantly differentially regulated^53^. Genes were ranked by their Wald statistic and their log2 fold-change values and used as input for gene set enrichment analysis (GSEA) on the molecular signatures database (MSigDB)^54–57^. Any signature with adjusted p-value < 0.01 was taken as significant. List of differentially expressed genes and significantly enriched pathways are reported in **Supplementary Table 1**.

Differential isoform usage was estimated with IsoformSwitchAnalyzeR^58^. Briefly, Salmon isoform count matrices from every sample were imported using *importRdata*^59^, using Gencode v33 exon annotations and nucleotide sequences. Single-isoform genes were filtered, as well as any genes below an expression cutoff of 1. Isoform switch testing was performed using *isoformSwitchTestDEXSeq* with default parameters^60,61^. All tissue types were tested separately, with COVID versus Control as the experimental condition. Differentially spliced genes were annotated as such: coding potential was assigned using CPAT^62^, protein domains were predicted with Pfam^63^, signal peptides were predicted with SignalP^64^, and intrinsically disordered regions were predicted using IUPred2a^65^, all using default parameters. Final isoform switch lists were generated using a differential isoform fraction cutoff of 0.1, a coding cutoff of 0.725, and non-coding ORFs removed. Gains and losses of predicted splicing events were predicted with *extractSwitchingSummary* and the functional significance of events was predicted with *extractConsequenceSummary*. Finally, differentially spliced pathways were estimated using PathwaySplice^19^, a gene feature-aware pathway analysis tool, using no GO size limitation, the *Wallenius* method of hypergeometric bias adjustment, and an exon-counting bin size of 20. Pathways were annotated against the Reactome Pathway Database^66^.

### TCEM Analysis

We used the method described by Danko et al^67^ to identify potential T Cell Expressed Motifs (TCEMs) in transcriptomic data from this study. Briefly, that method includes assembling IgH sequences using MiXCR^68^, then selecting 5 amino acid k-mers from the CDR3 regions of these sequences using three specific patterns specified by Bremel et al^69^. The whole set of 5aa sequences, with abundances, is the TCEM repertoire for a sample. We normalized the abundances of our TCEM repertoires by the total number of TCEMs of each pattern in a sample to yield relative abundances.

We mapped the TCEM repertoire from each sample to canonical SARS-CoV-2 protein sequences. We took every 5 amino acid subsequence from each protein and matched these sequences to the 5aa sequences in our TCEM repertoire. Any 5aa sequence which did not uniquely identify a single SARS-CoV-2 protein was discarded. We mapped each remaining 5aa sequence to a position on the SARS-CoV-2 genome by offsetting the position within the protein to the position by the position of the protein on the genome. Some 5aa sequences occurred multiple times in the same protein. For these sequences we arbitrarily used the highest (most 3’) coordinate.

## Extended Data Figures

**Extended Data Figure 1. Hierarchical clustering of viral genes in the lung.**

**a.** ERCC and HK normalized nCounter gene expression values were centered and rescaled. The resulting Z-scores are shown in this unsupervised clustering. COVID-19 positivity, where applicable, is indicated above columns. **b.** Summary of days of disease durations of each high/low groups.

**Extended Data Figure 2. Differentially expressed genes across three conditions.**

**a.** Ternary plot for Flu vs COVID-19 high vs COVID-19 low, **b.** Ternary plot for Non-viral ARDS vs COVID-19 high vs COVID-19 low.

**Extended Data Figure 3. Deconvoluted cell type proportions by conditions.**

From GeoMx data, cell proportions for each ROIs were plotted to compare between COVID-19 vs. normal control. The median and quartiles are noted by the box plot inside. P-value two-tailed t-tests were done to compare the means (ns: non-significant, *:p <= 0.05, **: p <= 0.01, ***: p <= 0.001, and ****: p <= 0.0001).

**Extended Data Figure 4. Examples of Trichrome stained lung tissue regions used for quantification.**

**a-d.** COVID-19 high cases: COVID1, COVID2, 55 and 59 showing mild increase in collagen zones with focally increased spindled cells, best seen in (b). **e-h.** COVID-19 low cases: COVID 17, 21, 73 and 86 with marked areas of increased collagen containing spindled cells, best seen in (**f,g**). **i-k.** normal lungs: 4, 103 and 104 showing blue staining collagen without expanded zones of cellularity (Masson trichrome, for all panels original magnification × 50). **l.** Quantification of collagen-rich fibroblastic zones, *p-value<0.01.

**Extended Data Figure 5. Quantitative descriptions of correlations and cell-type counts entropy values.**

**a.** Quantitative, pairwise comparison of the cell type correlations. To quantitatively compare correlation patterns in Figure 3a, the correlations are compared pairwise by Fisher’s test. The transformed scores and p-values are shown. **b.** Entropy calculations for the cell type counts. Entropy of cell type counts are visualized by tissue types, as in Figure 3c. Area marked in darker grey indicates significantly insignificant results.

**Extended Data Figure 6. ssGSEA scores by conditions.**

The single-sample enrichment scores of **a.** macrophage, **b.** neutrophil, and **c.** T and NK cell related gene sets were evaluated. The scores were averaged and compared across COVID-19 high and low, ARDS, flu, relative to normal.

**Extended Data Figure 7. Viral Genome Reconstruction from Total RNA-Seq Data.**

**a.** For each patient sample per tissue, the percentage of Total RNA-seq reads mapped uniquely to SARS-CoV-2 and no other taxon is shown, against the percentage of the viral genome (Wuhan reference) that was covered with ≥10 reads during genome assembly. **b.** Normalized expression value per gene body in the SARS-CoV-2 genome for samples where at least 10% of the viral genome was covered with ≥10 reads. **c.** Heatmap of variant alleles across the viral genome, grouped by patient.

**Extended Data Figure 8a. Representative images of lung morphology of the cases analyzed by GeoMx spatial transcriptomics.**

**a-h.** COVID cases (a) COVID1 (b) COVID2 (c) 55L (d) 59L (e) COVID 17 (f) COVID21 (g) 73L (h) 86L; **i-j.** ARDS and influenza (i) Flu1 (j) Flu2; **k-m.** ARDS non-influenza (k) ARDS1 (l) ARDS2 (m) ARDS3; **n-p.** Normal lung (n) NL4 (o) Archoi103 (p) Archoi 104. Hematoxylin and eosin stain, Original magnification × 100 except for c and d (x 50).

**Extended Data Figure 8b. Sample collection strategy for COVID autopsy samples from complete adult cases.**

Two cases with representative hematoxylin and eosin images are shown: **a-e.** Case 58 and **f-j.** Case 73. (a, f) Lung, (b,g) Kidney, (c,h) Heart, (d,i) Liver and (e,j) Mediastinal lymph node (Hematoxylin and eosin stain, (b,d,e,i,j) original magnification × 50, (a,c,f,g,h) original magnification × 100).

**Extended Data Figure 9. All dysregulated pathways across tissue types.**

Pathways found to be dysregulated based on isoform switch analysis are shown for each tissue type examined. Only pathways in which at least one tissue had log fold change > 1 and adjusted p-value < 0.05 were considered. Odds ratio is shown per pathway, and pathways are colored by adjusted p-values (with no shading for non-significant effects).

**Extended Data Figure 10. Correlation of different sample types and COVID-19 status.**

Correlations between each sample type (label) gene expression matrix and other tissue/sample types are shown as a correlation range (red, high up to 1.0 and blue, low down to −1.0). The p-values represent the significance of the correlation with * p-value < 0.05, ** p-value < 0.01, and *** p-value < 0.001.

