## Supplementary figures and images for "Systemic Tissue and Cellular Disruption from SARS-CoV-2 Infection revealed in COVID-19 Autopsies and Spatial Omics Tissue Maps"

### Extended Data Figure 1

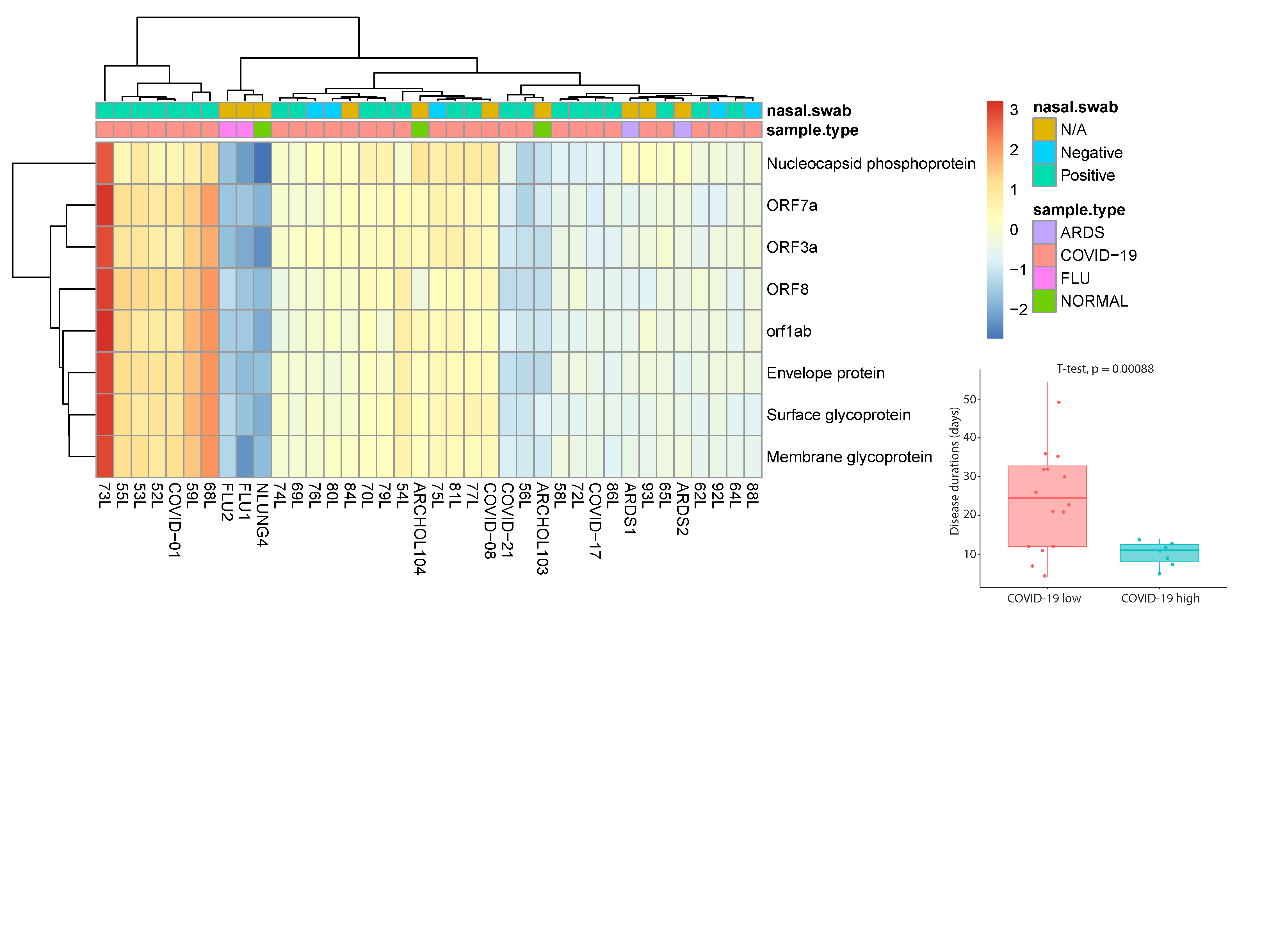

### Extended Data Figure 2

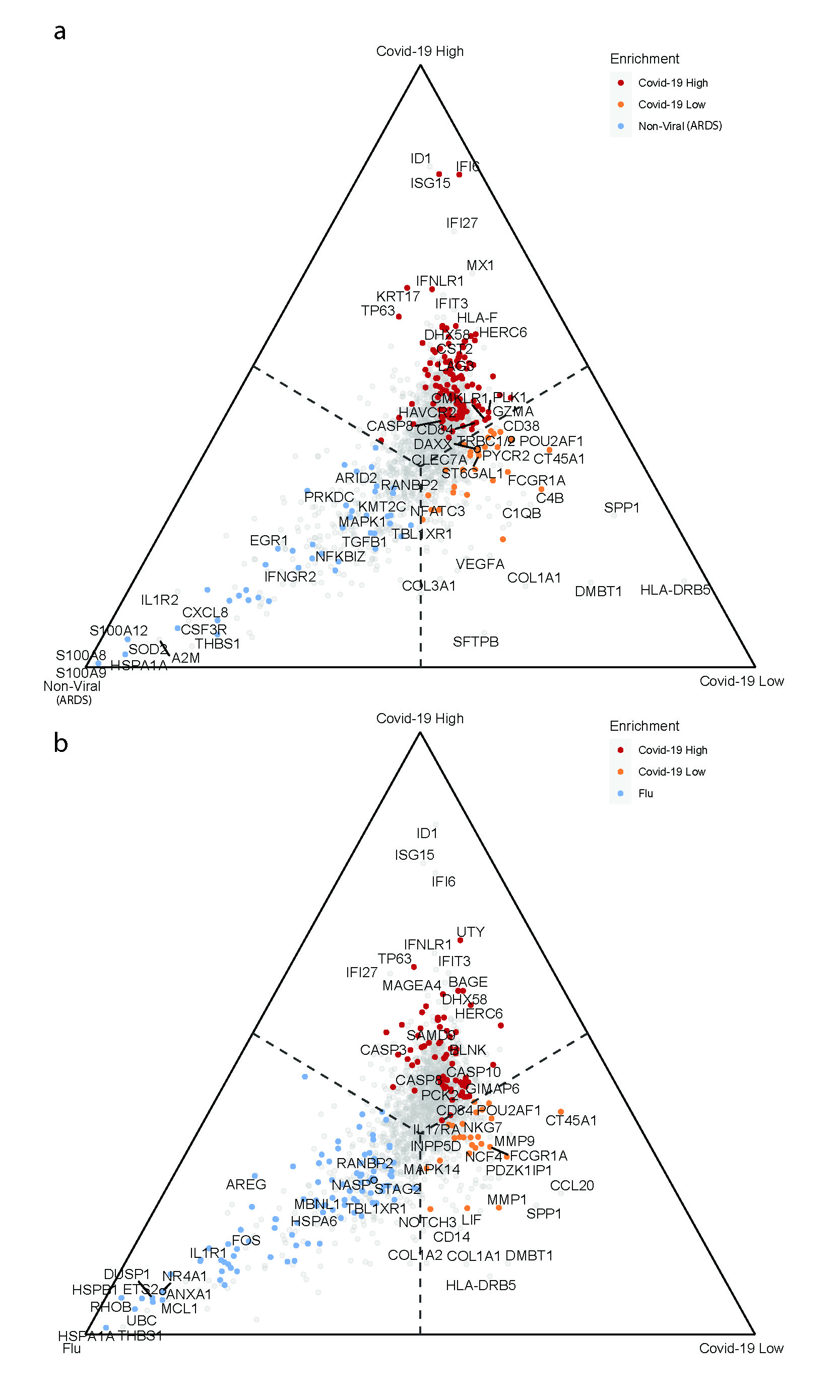

### Extended Data Figure 3

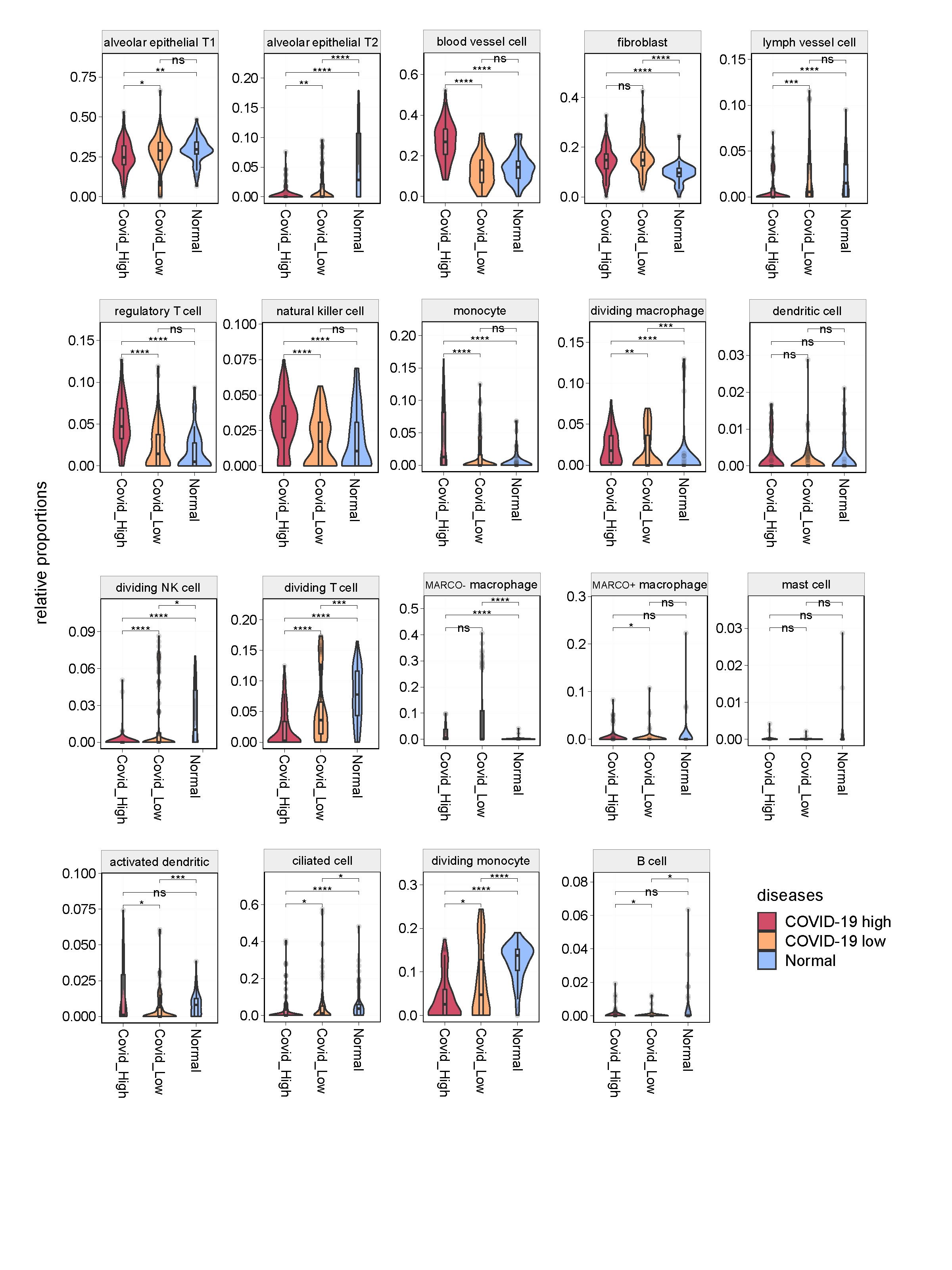

### Extended Data Figure 4

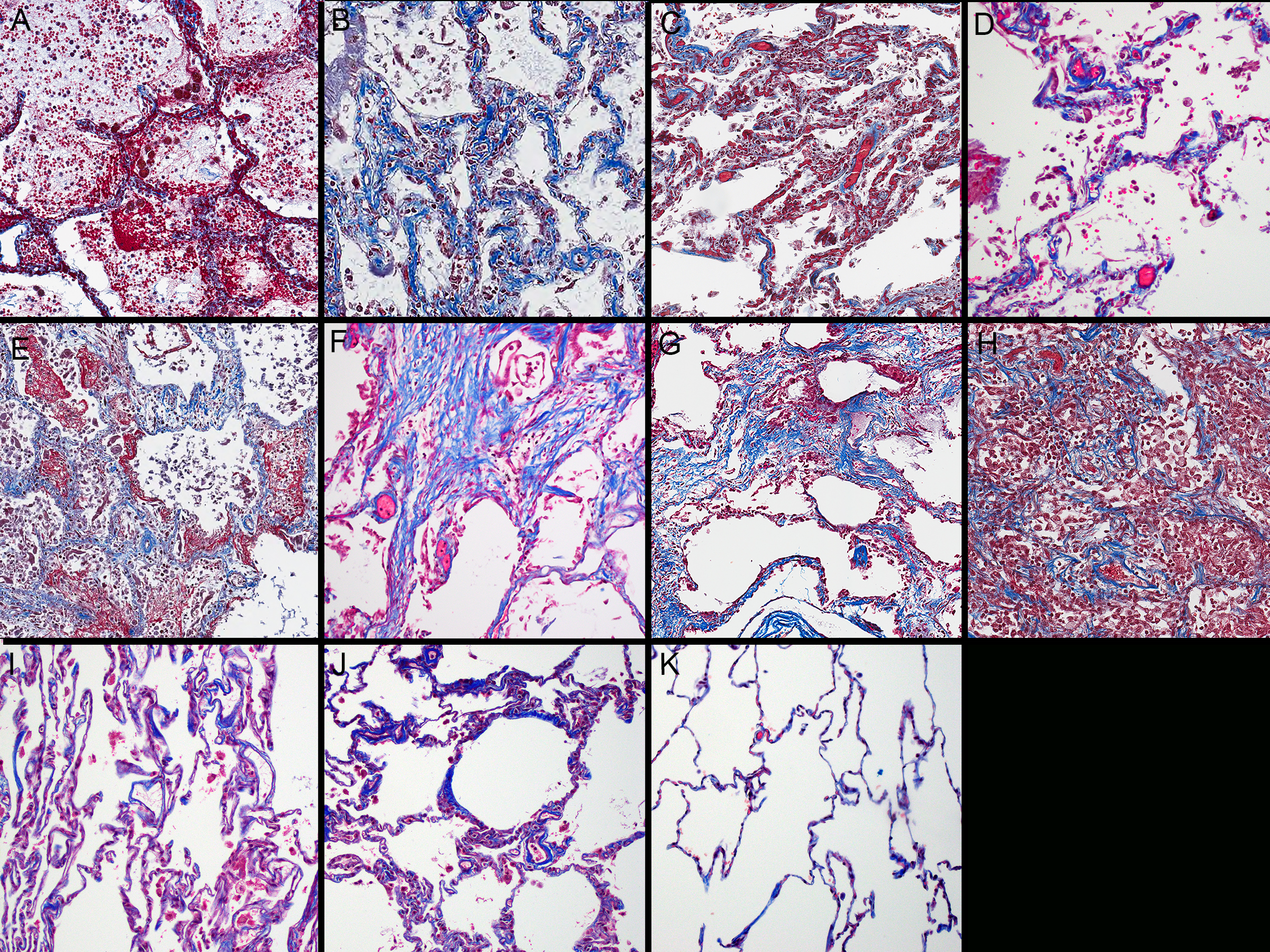

### Extended Data Figure 5

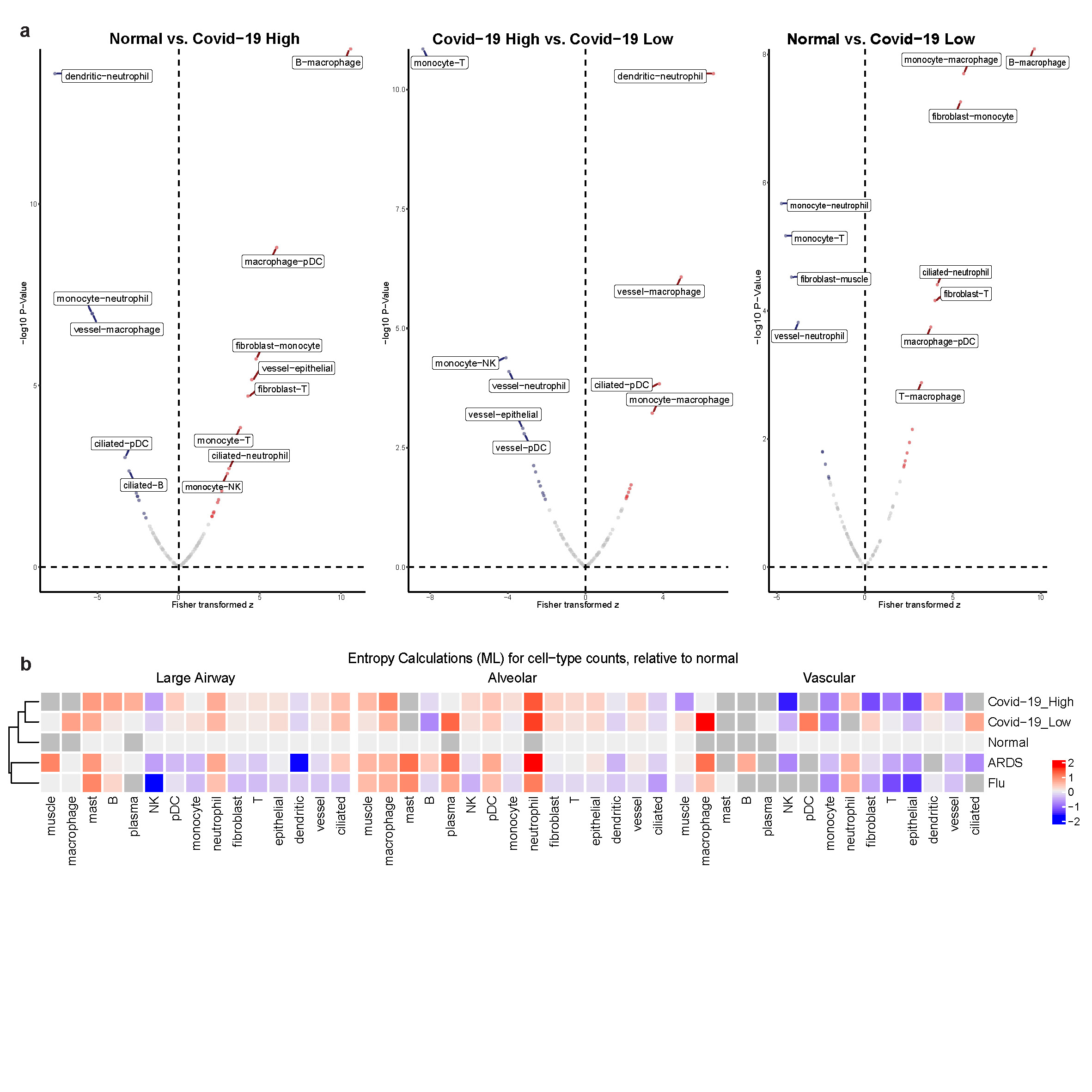

### Extended Data Figure 6

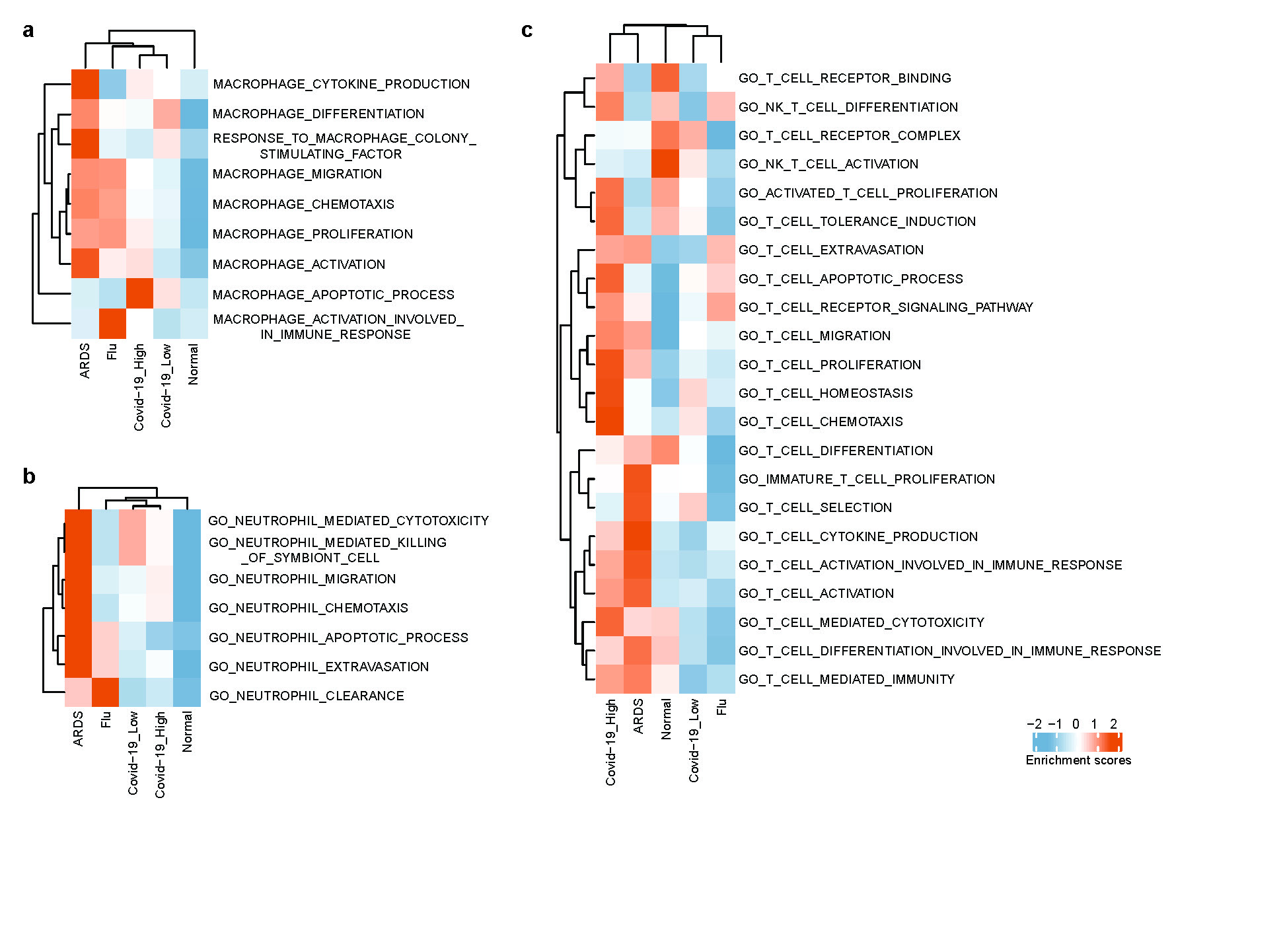

### Extended Data Figure 7

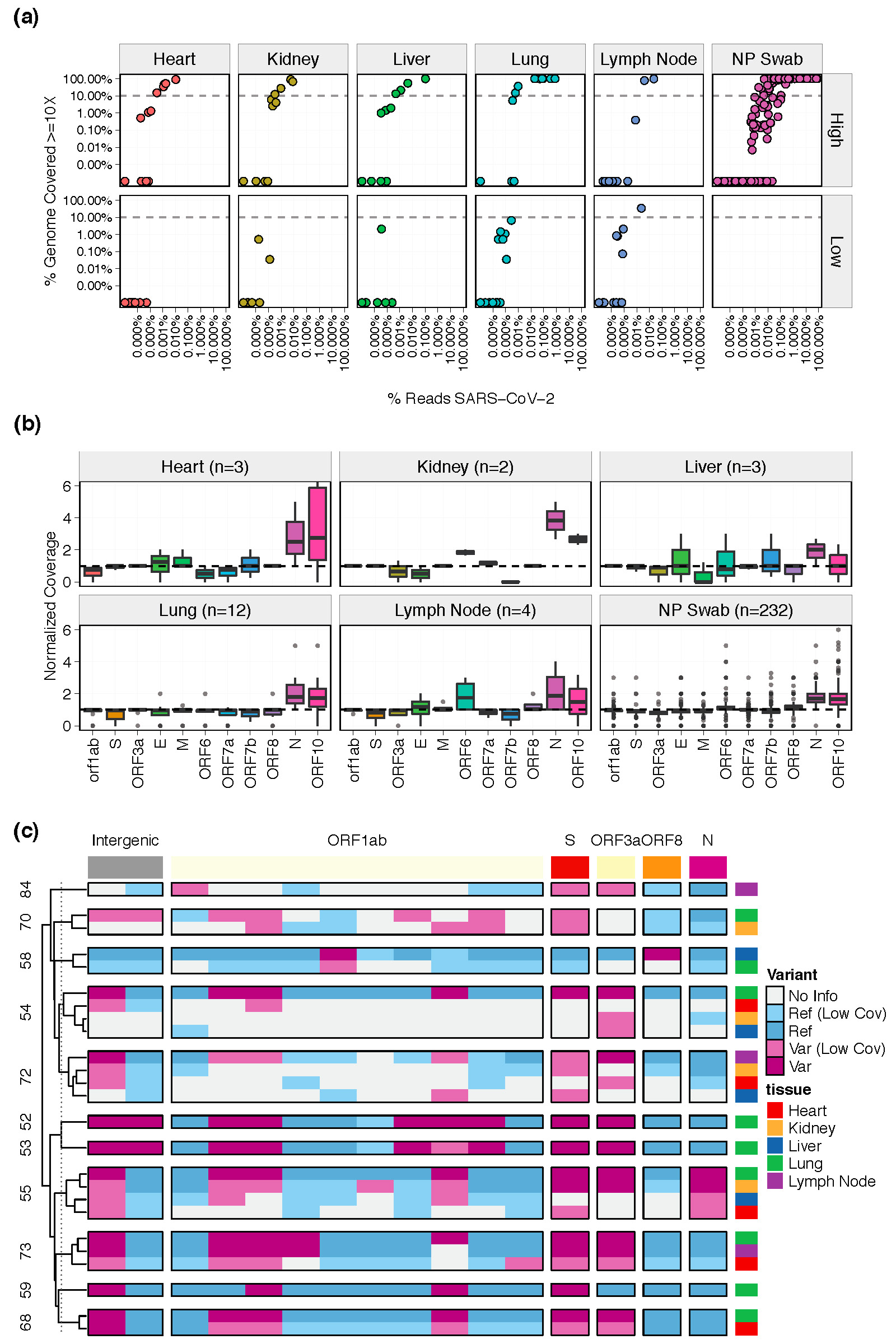

### Extended Data Figure 8a

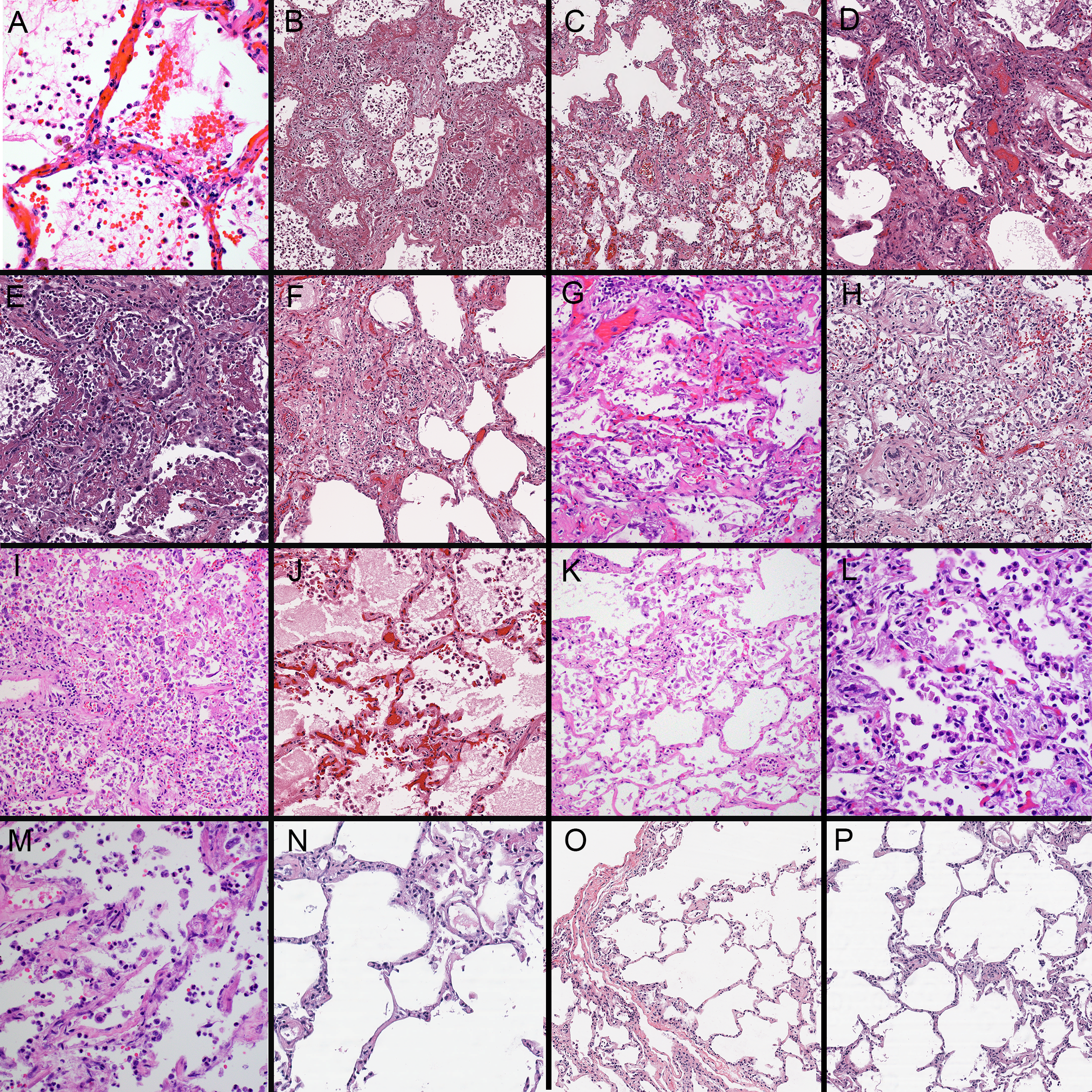

### Extended Data Figure 8b

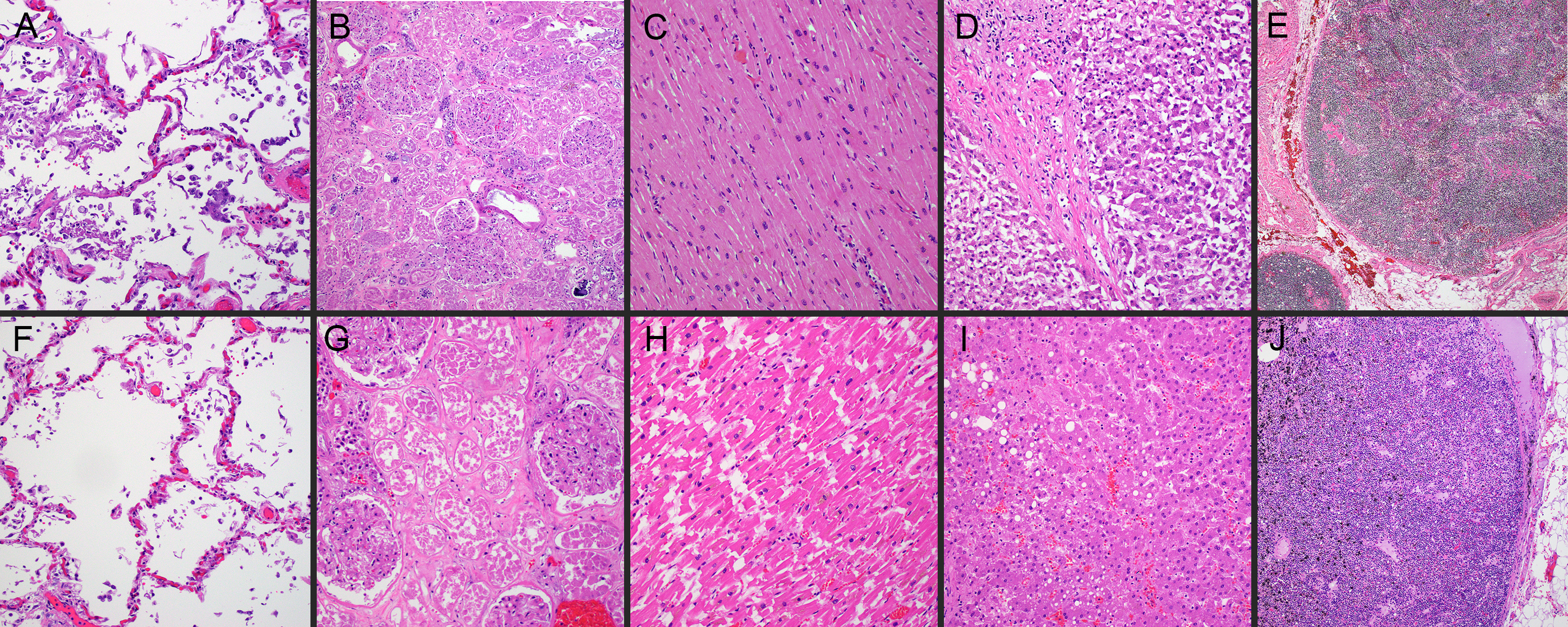

### Extended Data Figure 9

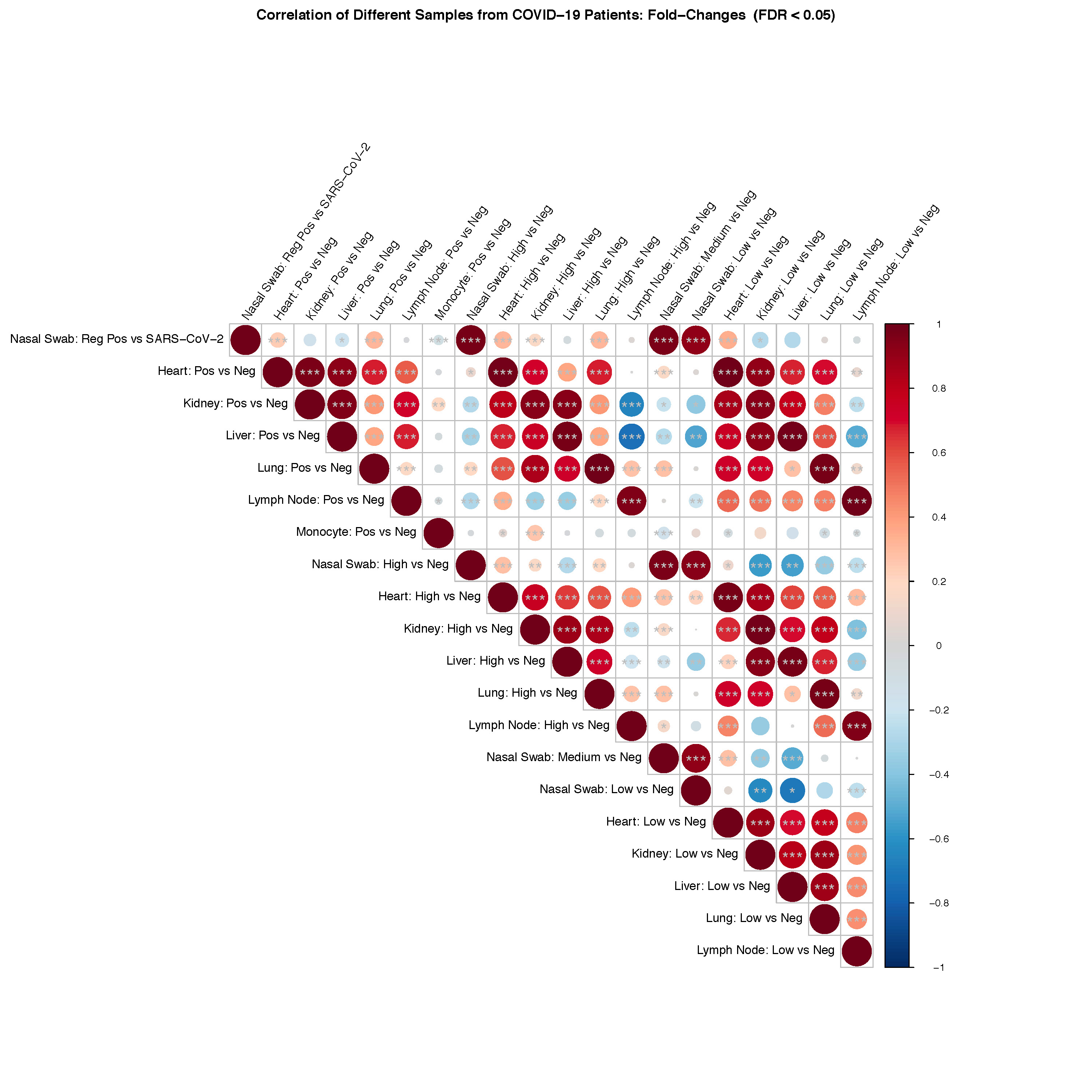

### Extended Data Figure 10

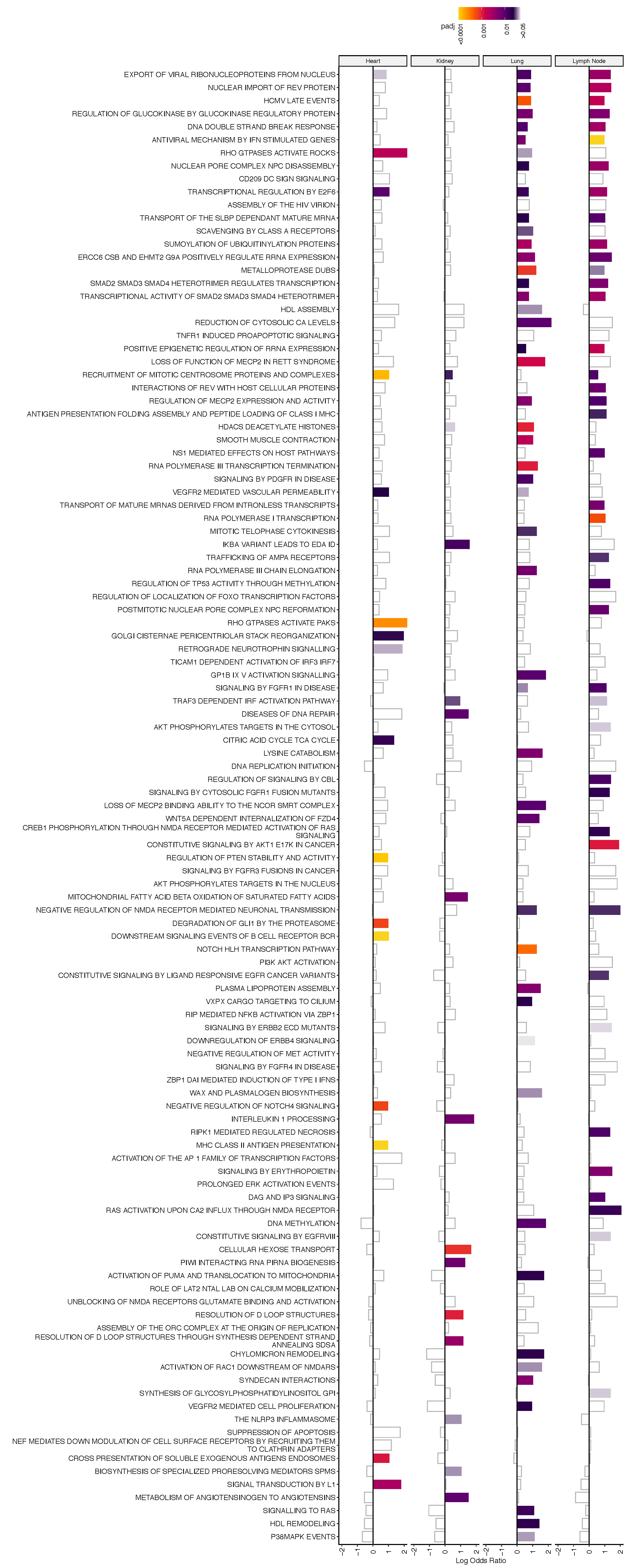
